# Structural basis of insulin receptor antagonism by bivalent site 1-site 2 ligands

**DOI:** 10.1101/2025.08.23.671589

**Authors:** Amber Vogel, Alan Blakely, Yuankun Dao, Nai-Pin Lin, Danny Chou, Christopher P. Hill

## Abstract

Congenital hyperinsulinism (HI) is a rare genetic disease characterized by overproduction of insulin. One class of potential HI treatments is insulin receptor (IR) antagonists like S961 and Ins-AC-S2, peptides composed of binding segments for each of the IR sites capable of binding insulin: site 1 and site 2. Notably, S597 – containing the same IR binding segments as S961 but in the opposite order (site 2-site 1) – is an IR agonist rather than an antagonist. Using cryo-EM, we show how both S961 and Ins-AC-S2 bind an inactive conformation of IR, thereby explaining their antagonism. Furthermore, our structures reveal how agonist vs. antagonist activity is dictated by the order of site 1- and site 2-binding modules in bivalent ligands. Additionally, we uncover subtle differences between the binding mechanisms of S961 and Ins-AC-S2 to IR, which include displacement or engagement of αCT, respectively, and a novel binding interface between the Ins-AC-S2 insulin and the receptor. These structural insights may inform development of next generation IR antagonists for treatment of HI.

## INTRODUCTION

Congenital hyperinsulinism (HI) is a rare genetic disease, occurring in roughly 1/25,000-1/50,000 infants, that is characterized by overproduction of insulin by pancreatic beta cells, leading to persistent hypoglycemia^1,2^. Left untreated, patients with HI can experience seizures, brain damage, learning disabilities, and death. HI can be caused by a vast number of genetic mutations and other transient causes and metabolic diseases, making treatment complicated and case-specific^3^. Most available treatments target endogenous insulin secretion pathways but are not universally effective, carry serious side effects, and drug resistance can develop^3^. This forces many patients to undergo surgical removal of part or most of the pancreas, which can result in type 1 diabetes. Thus, new treatment options for patients with HI are desirable therapeutic targets.

Insulin signaling is mediated by binding of insulin to the insulin receptor (IR), a member of the receptor tyrosine kinase family that is critical for import of glucose into the cell^4^. In the absence of insulin, the fibronectin stalks that point to the cell membrane are separated and hold the intracellular tyrosine kinase (TK) domains apart, preventing trans-autophosphorylation of the TK domains required for downstream signaling (Fig. 1a). Binding of insulin to the ectodomain (ECD) results in dramatic conformational changes that bring the fibronectin stalks together, allowing for receptor activation (Fig. 1b)^4,5^. IR is a disulfide crosslinked homodimer, where each protomer contains α and β chains comprising multiple domains (Fig. 1c). Each α chain contains two binding sites for insulin: site 1, composed of the leucine-rich 1 (L1) domain and the alpha C-terminal helix (αCT) of the other protomer; and site 2, composed of the fibronectin type-III 1 (FnIII-1) domain. Due to its dimeric structure, there are four binding sites on IR: site 1 and site 2 on one receptor half, and the corresponding site 1’ and site 2’ of the other receptor half. Differential splicing of the IR gene results in A and B isoforms, which differ by the absence or presence of an 12-residue extension at the C-terminus of the α chain^6^. A crystal structure shows how apo/inactive IR adopts an inverted-V conformation with the extracellular fibronectin stalks separated by ∼120 Å (Fig. 1d)^7^. Multiple cryo-EM structures show activated complexes with 1:1, 1:2, and 1:3 IR:insulin stoichiometries^8–12^. Interestingly, the single insulin of a 1:1 complex may be sufficient for activation, as demonstrated by asymmetric structures showing close proximity of the fibronectin stalks (Fig. 1e)^8,12^.

**Figure 1:**
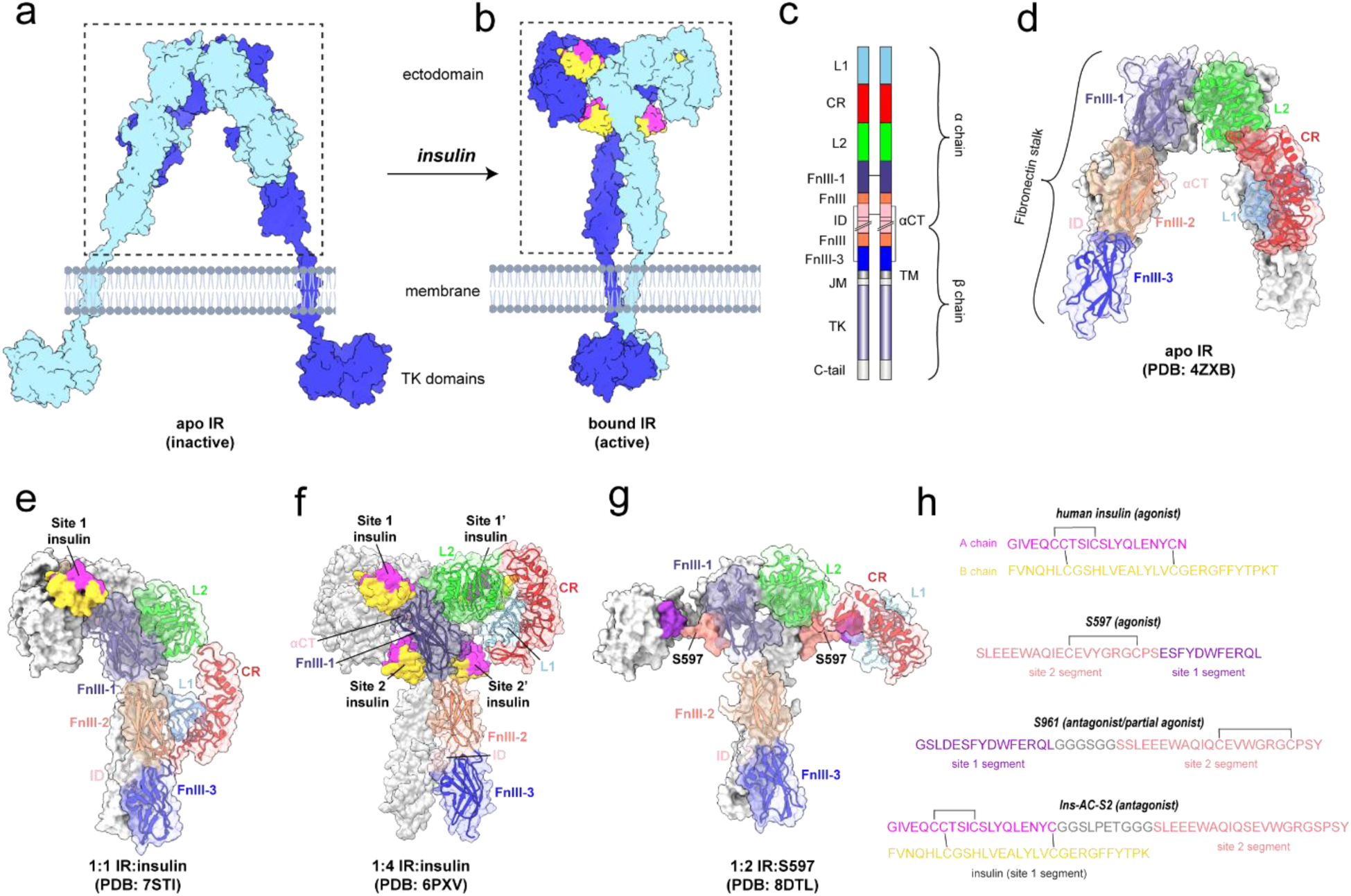
Insulin receptor and ligand structure and domain organization. (a-b) Cartoon of (a) the apo/inactive FL-IR illustrating the physical separation of the TK domains, and (b) the insulin-bound/active FL-IR showing dimerization of the TK domains. Cartoon generated using PDBs: 6PXV, 4ZXB, Alphafold3^43^, and BioRender.com. The ectodomain (ECD) is boxed in (a-b). (c) Domain organization and disulfide linkages of FL-IR drawn to scale using illustrator of biological sequences^44^. The α and β chains are indicated and both protomers are shown. (d) Structure of the apo IR ECD in the inverted-V conformation (PDB: 4ZXB). (e) Structure of the IR showing only the ECD bound to one site 1 insulin in an asymmetric, active conformation (PDB: 7STI). (f) Structure of the IR showing only the ECD bound to four insulin molecules in the activated T-shape conformation (PDB: 6PXV). (g) Structure of the IR showing only the ECD bound to two S597 molecules (PDB: 8DTL), where IR adopts a variation of the active T shape conformation. For structures shown in panels (d-g), one IR protomer is shown in grey and the other color coded by domain according to the color scheme shown in panel (c): leucine-rich 1 domain (L1, light blue), cysteine-rich domain (CR, red), leucine-rich 2 domain (L2, green), fibronectin type-III 1 domain (FnIII-1, dark purple), fibronectin type-III 2 domain (FnIII-2, coral), alpha C-terminal helix (αCT, pink), insert domain (ID, pink), and fibronectin type-III 3 domain (FnIII-3, dark blue). (h) Sequences of human insulin and IR ligands used in this study with site 1- and site 2-binding segments indicated. Human insulin A and B chains are colored magenta and gold, respectively. S597 and S961 site 1 and site 2 segments are shown in purple and salmon, respectively. The S961 flexible linker is shown in grey. The Ins-AC-S2 A chain contains binding segments for site 1 and site 2 (colored in magenta and salmon, respectively), with the linker shown in grey. The Ins-AC-S2 B chain is shown in gold. Disulfide bonds are indicated.

Antagonists that bind IR and inhibit activation by insulin may have potential to treat HI. Known antagonists include the 43-residue peptide S961^13^, the monoclonal antibody IRAB-B^14^, and the insulin derivative Ins-AC-S2^15^. S961 comprises a site 1-binding peptide, a short linker, and a site 2-binding peptide (Fig. 1h)^16^. The site 1-binding peptide by itself is an IR agonist with an affinity of 11-40 nM^16,17^. S661, the C-terminal acid form of S961 with otherwise identical properties^13^, has reported IC_50_ values of 2.4 nM and 7.4 nM for human IR-A and -B over-expressing HEK293 cells, respectively^18^. Remarkably, the same binding segments in the reverse order (site 2-site 1) produced an IR agonist called S597 (Fig. 1h), indicating that the organization of binding segments is critical in determining whether an effector functions as an agonist or an antagonist. Structures are known for the isolated site 1-binding peptide complexed with the “microreceptor” construct that comprises just the L1, CR, and αCT domains,^17^ and of the site 2-binding peptide bound to the full-length insulin receptor^19^. Furthermore, the structure of full-length IR in complex with S597 showed IR in an active T-shape conformation with close proximity of the FnIII stalks (Fig. 1g)^19^.

Unfortunately, several pitfalls prohibit the use S961 as a therapeutic for HI. First, S961 displays partial agonism at low concentrations^20^. Second, S961 exhibits undesirable mitogenic signaling at low concentrations (1-10 nM) in L6-hIR, MCF-7, and CHO-hIR cells. Although S961 is not suitable as a therapeutic, it inspired the recent development of Ins-AC-S2, which displayed an IC_50_ of 3.0 nM^15^. Ins-AC-S2 comprises an insulin molecule ligated at the A chain C-terminus to the site 2-binding segment of S961 with two Cys to Ser point mutations (Fig. 1h). Importantly, unlike S961^20^, Ins-AC-S2 does not display partial agonism at low concentrations^15^.

To better understand how S961 and Ins-AC-S2 antagonize IR, we determined their cryo-EM structures in complex with IR. Both antagonists bind an inverted-V conformation of IR with wide separation of the fibronectin stalks, explaining their antagonistic activity and showing how the antagonist site 1-site 2 orientation stabilizes the inactive state. Intriguingly, we were able to identify two unexpected but important structural features for antagonism. First, the site 1-binding segment of S961 structurally replaces αCT, a receptor helix involved in insulin binding and negative cooperativity. Consistent with direct competition, we found that S961 binds stably to one receptor half but undergoes dynamic association and dissociation from the other receptor half that may be detrimental for forming a stably inhibited complex with S961, even at micromolar concentrations. Second, we discovered a novel binding interface between Ins-AC-S2 and the IR FnIII-2 and insert domains, which we call site 1Ʌ. This interface contributes to the antagonism of Ins-AC-S2, is only present in the inactive conformation, and may shed light on the initial stages of native insulin binding to the receptor during canonical IR activation. These findings can inform the development of next generation antagonists with improved properties for the treatment of HI.

## RESULTS AND DISCUSSION

### Structure of the IR/S961 complex

To visualize S961 bound to IR, two IR-A constructs were developed: (i) an ectodomain construct (residues 1-930, referred to as ECD) with C-terminal 6x-His and 2x-FLAG tags, and (ii) full-length IR (residues 1-1343, referred to as FL-IR) with a C-terminal 2x-FLAG tag. Both proteins were expressed in HEK293/17 SF cells and purified either from the growth media (ECD) or cell membranes (FL-IR). ECD and FL-IR were each incubated with S961 and applied to Quantifoil holey carbon or self-wicking nanowire grids. Movies were collected on a Titan Krios equipped with a Gatan K3 camera and energy filter. 3D reconstructions were first performed on particles obtained from FL-IR and ECD separately, resulting in a 3.88 Å map from 497,687 particles (FL-IR/S961) and a 3.93 Å map from 248,634 particles (ECD/S961). Subsequently, particles from datasets obtained for FL-IR and ECD bound to S961 were combined and resulted in a 3.68 Å reconstruction from 378,182 particles (FL-IR+ECD/S961, Fig. 2a-b). Since no apparent differences were observed between the FL-IR/S961 and ECD/S961 maps (Sup. Fig. 4a-c), and the resolution improved when particles from all datasets were combined, subsequent structural analyses focused on reconstructions obtained from the combined data.

**Figure 2:**
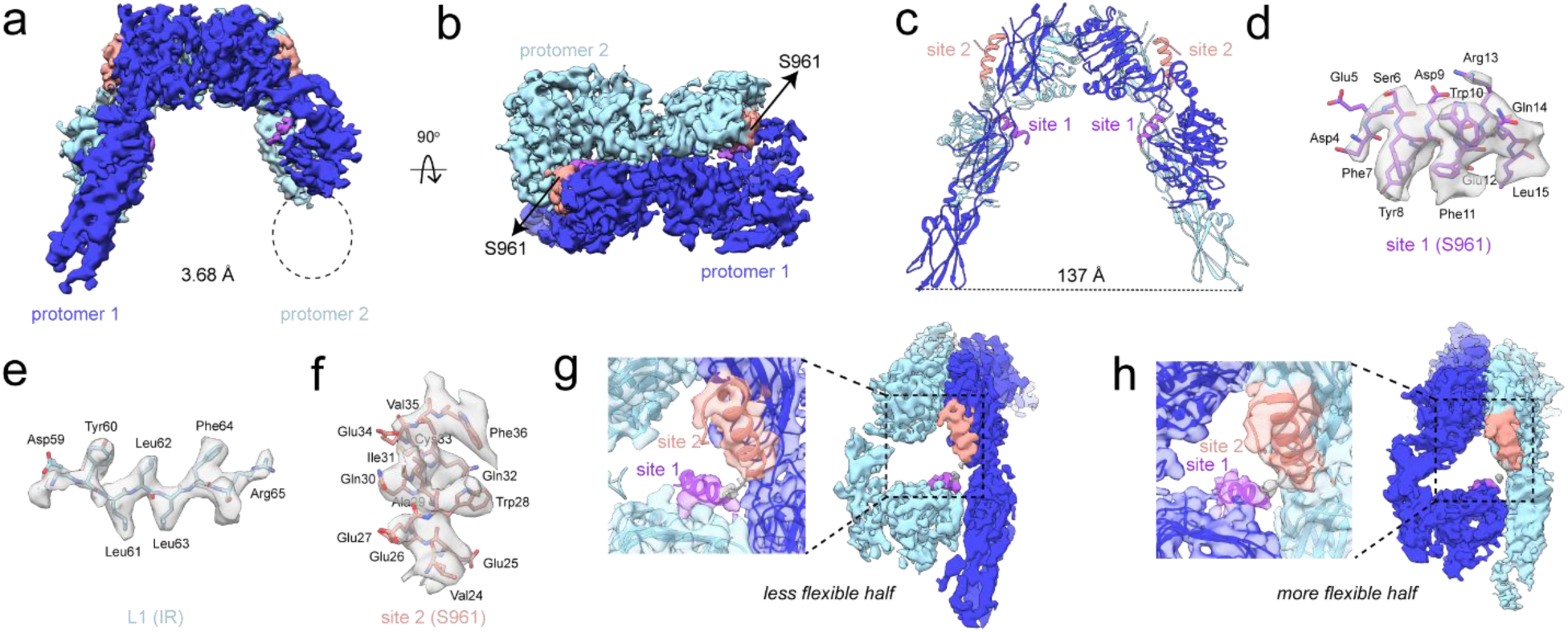
Structure of IR bound to S961. (a-b) Cryo-EM density map determined to 3.68 Å from 378,182 particles showing (a) front view and (b) top view of the receptor. Protomers 1 and 2 are colored in dark and light blue, respectively. The density for S961 site 1- and site 2-binding segments are colored in purple and salmon, respectively, and the linker density is shown in grey. The region with missing density corresponding to the FnIII-3 domain of protomer 2 is circled. (c) Model of the IR/S961 complex showing the separation distance of the fibronectin stalks (distance measured from L909 to L909’ on the opposite protomer). (d-f) Representative map and model of (d) S961 site 1 helix, (e) β-strand from L1 belonging to IR site 1, and (f) S961 site 2 helix. (g-h) Side views of IR/S961 density map and model from (a) showing (g) the better reconstructed, and (h) the less well reconstructed receptor halves with contour levels adjusted to see S961 at both sets of binding sites. Zoomed in views of the map and model overlayed at both sets of binding sites are shown with binding sites labeled.

IR bound to S961 adopted an inverted-V conformation, consistent with S961 being an antagonist (Fig. 2a-c). The density map was sufficient to model most domains of the receptor, and adequate side chain density was observed at both site 1 and site 2, providing confident helical register of S961 (Fig. 2d-f). However, map resolution was strikingly asymmetric between the two receptor halves (Fig. 2a-b, Sup. Fig. 4c). On the more clearly reconstructed half, the full fibronectin stalk was visualized, while the less clearly reconstructed half lacked density for the FnIII-3 domain, indicating a high degree of conformational heterogeneity. However, S961 was clearly visualized bound to both receptor halves, indicating an IR dimer:S961 stoichiometry of 1:2 for the majority of particles (Fig. 2g-h).

### Ligand binding site organization and agonism vs. antagonism

To better understand the mechanism of S961 antagonism, we compared our model to several previously determined structures. Compared to structures of IR complexed with isolated site 1 or site 2 components of S961^17,19^, we found no significant differences in peptide coordination relative to our full-length S961 complex structure, establishing that the site 1- and site 2-binding helices consistently bind IR at the same sites and in the same orientation (Fig. 3a-b). Due to the missing density for the FnIII-3 domain of one stalk, it was difficult to determine an exact separation distance between the fibronectin stalks in the IR/S961 complex. However, by rigid body fitting of the entire fibronectin stalk, we determined an approximate stalk separation distance of 137 Å (Fig. 2c). This is similar to the fibronectin stalk separation distance in the apo IR structure (112 Å)^7^. Furthermore, the conformations of the L1/CR/L2 domains of each protomer relative to the adjacent fibronectin stalks were nearly identical between the apo structure and IR/S961 complex (Fig. 3c), indicating that any difference in fibronectin stalk distance is the result of rigid body hinging between the two receptor halves.

**Figure 3:**
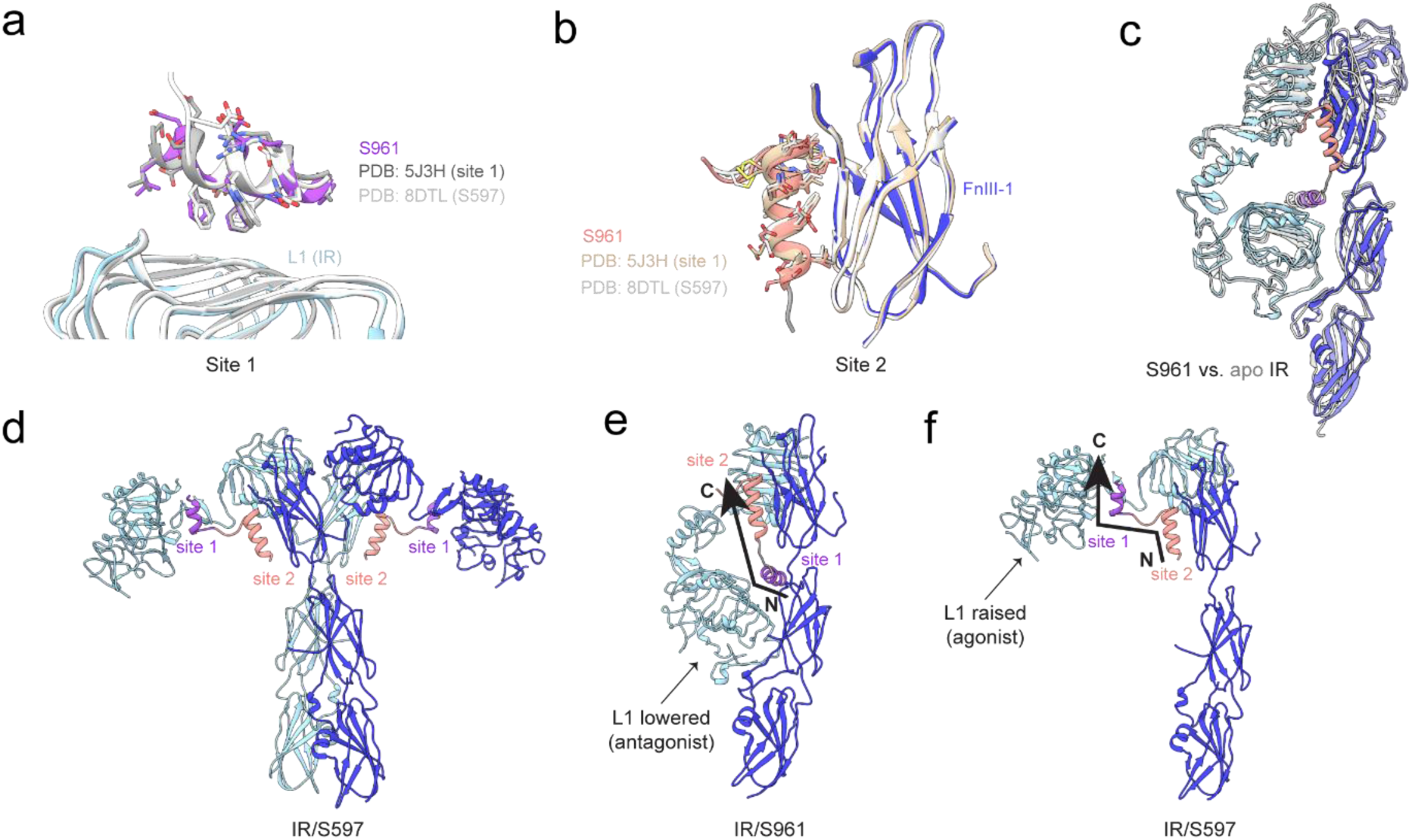
Structural comparison of the IR/S961 complex and isolated binding segments, apo IR, and the IR/S597 complex. (a) IR/S961 model showing the site 1 helix overlayed with S519C16 (site 1 component, grey, PDB: 5J3H) and S597 (white, PDB: 8DTL) aligned to L1. (b) IR/S961 model showing the site 2 helix overlayed with isolated site 2 peptide (tan, PDB: 8DTM) and S597 (grey, PDB: 8DTL) aligned to FnIII-1. (c) Side view showing half of the IR/S961 complex (color) overlayed with apo IR (white, PDB: 4ZXB). (d) Published structure of the IR/S597 complex (PDB: 8DTL). (e-f) Side view showing one half of (e) the IR/S961 complex and (f) the IR/S597 complex highlighting the N- to C-terminal orientation of S961 and S597 with arrows. Site 1 and site 2 are labeled in purple and salmon, respectively.

A structure of mouse FL-IR in complex with S597, a 31-residue peptide agonist of the IR that contains the same site 1- and site 2-binding segments as S961 but in the opposite order (Fig. 1g-h) adopts an activated T-shape conformation (Fig. 3d)^19,21^. Interestingly, the site 1- and site 2-binding segments of S597 each individually bind their sites on IR in essentially the same way as in the inactive inverted-V complex with S961, with the only difference being one less helical turn of the S597 site 1 helix due to the linker connecting sites 1 and 2 (Fig. 3a-b). Overall, this comparison further supports the conclusion that whether bivalent ligands are agonists or antagonists can be dictated simply by the order of site 1- and site 2-binding segments.

To understand how binding site organization dictates agonist versus antagonist activity, we compared our IR/S961 structure to the published IR/S597 structure. In both cases, the peptide N-terminus points down toward the membrane with the C-terminus opposing. For S961, the N-terminal site 1-binding segment is membrane-proximal to the site 2 binding segment, forcing the L1 and FnIII-1 domains to be in similar relative positions as the apo conformation and incompatible with activated conformations (Fig. 3e). In contrast, for S597 the site 1-binding segment is parallel to the site 2-binding segment with respect to the membrane, allowing the L1 domain to adopt a raised position that is compatible with an active conformation (Fig. 3f). Thus, agonist/antagonist activity can be determined by bivalent ligands stabilizing the relative positions of the L1 (site 1) and FnIII-1 (site 2) domains.

### Conformational heterogeneity of the IR/S961 complex

Local resolution was strikingly asymmetric between the two receptor halves of the IR/S961 complex (Fig. 2a-b, Sup. Fig. 4c). We therefore performed local refinement separately on each half. Even after local refinement, one half was far better resolved than the other, indicating that motion originates from domain movement within the less resolved half, rather than rigid body movement of one receptor half relative to the other (Sup. Fig. 4j-k). We subsequently performed 3D variability analysis (3DVA), which indicated the well-resolved half of the complex was the anchor for particle alignment (Fig. 4a), while the L1/CR module of the other half of the receptor deviates from a perfectly symmetric structure and adopts a range of conformations. Motion originates from hinging between the CR and L2 domains that asymmetrically raises one of the L1/CR modules containing site 1. At the most symmetric end of the trajectory, the L1/CR module is in its lowest sampled position, strong density is apparent for S961 at site 1, and connecting density is prominent for the linker connecting sites 1 and 2, suggesting high occupancy of S961 (Fig. 4b). In contrast, when the L1/CR module is raised, density for S961 at site 1 is weaker and density is absent for the linker, suggesting lower occupancy or dynamics at site 1 (Fig. 4c). In all conformations, density is missing for FnIII-3 nearest the mobile L1/CR domains. These data suggest S961 association/dissociation from site 1 on one receptor half is coupled to dynamics of the adjacent fibronectin stalk.

**Figure 4:**
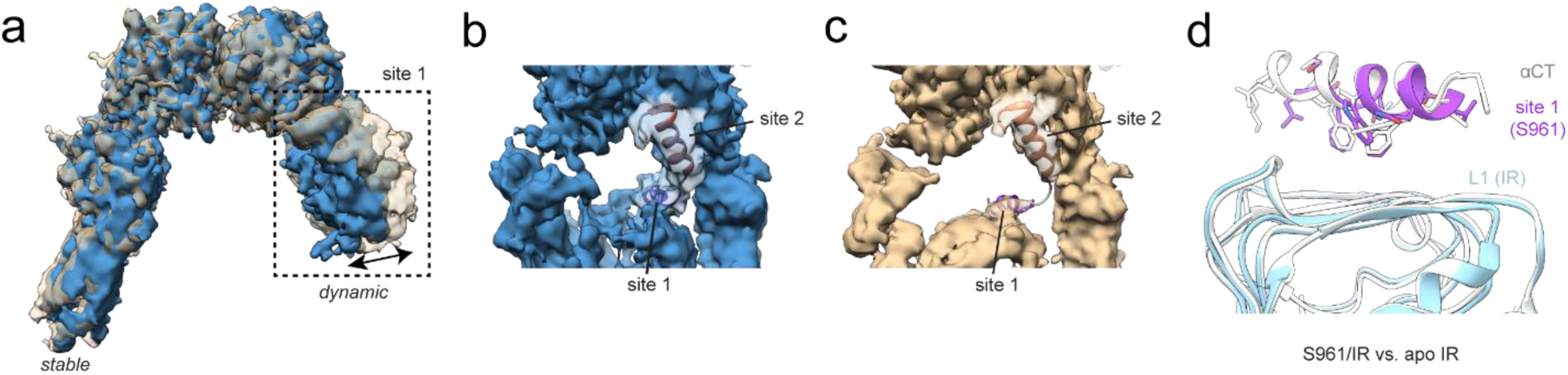
Competition for site 1 by S961 and αCT. (a) Overlay of density maps obtained from the two most different subsets of particles obtained by 3DVA shown in tan and steel blue. The dynamic receptor arm containing site 1 is boxed. (b-c) Zoomed in view of S961 at site 1 and site 2 fit into the density maps shown in (a). (b) Map showing strong density for S961 at site 1 determined to 3.99 Å resolution from 343,705 particles. (c) Map showing weak density for S961 at site 1 determined to 3.97 Å from 386,871 particles. Site 1 and site 2 helices of S961 are labeled. (d) IR/S961 site 1 helix overlayed with apo IR aCT (white, PDB: 4ZXB) showing sidechains pointed towards the L1 domain and aligned to the L1 domain.

S961 is reported to be a partial agonist at low concentrations, but the molecular basis for this phenomenon remains unknown^20^. Although we did not obtain a reconstruction of the IR in an active conformation, the mobility observed for the fibronectin stalk nearest the labile L1/CR module opens the possibility of transient sampling of active conformations. Since stalk motion is correlated with lesser density for the helical element (presumably S961) at site 1, we compared our IR/S961 model to apo IR (PDB: 4ZXB^7^), focusing on site 1 to understand the impact of S961/αCT dissociation. As seen in the structure of the site 1 segment of S961/S597 alone in complex with the receptor^17^, αCT and the S961 site 1 helix bind L1 in identical orientations and share similar binding interface residues, and thus are competitive binders of L1 (Fig. 4d). The weaker density at site 1 of one receptor half may reflect structural heterogeneity resulting from this competition. Competition between S961 and αCT likely destabilizes the antagonized IR/S961 complex, which may contribute to the partial agonism of S961 at low concentrations.

### Structure of the ECD/Ins-AC-S2 complex

We next sought to determine the cryo-EM structure of FL-IR bound to Ins-AC-S2, a potent, full antagonist^15^ that has a similar low nanomolar IC_50_ value as S961^18^ but slightly higher potency in our side-by-side antagonism assay (Sup. Fig. 5, Sup. Table 2). Ins-AC-S2 comprises human insulin fused to the site 2-binding component of S961/S597 with two Cys to Ser mutations (Fig. 1h). Sample preparation and cryo-EM data collection were performed as for the IR/S961 complex. The data quality improved when using the ECD construct rather than FL-IR, which ultimately produced a 3.64 Å map (Fig. 5a-b). The FL-IR and ECD complex maps displayed no significant differences, validating the use of the ECD for this structural analysis (Sup. Fig. 6).

**Figure 5:**
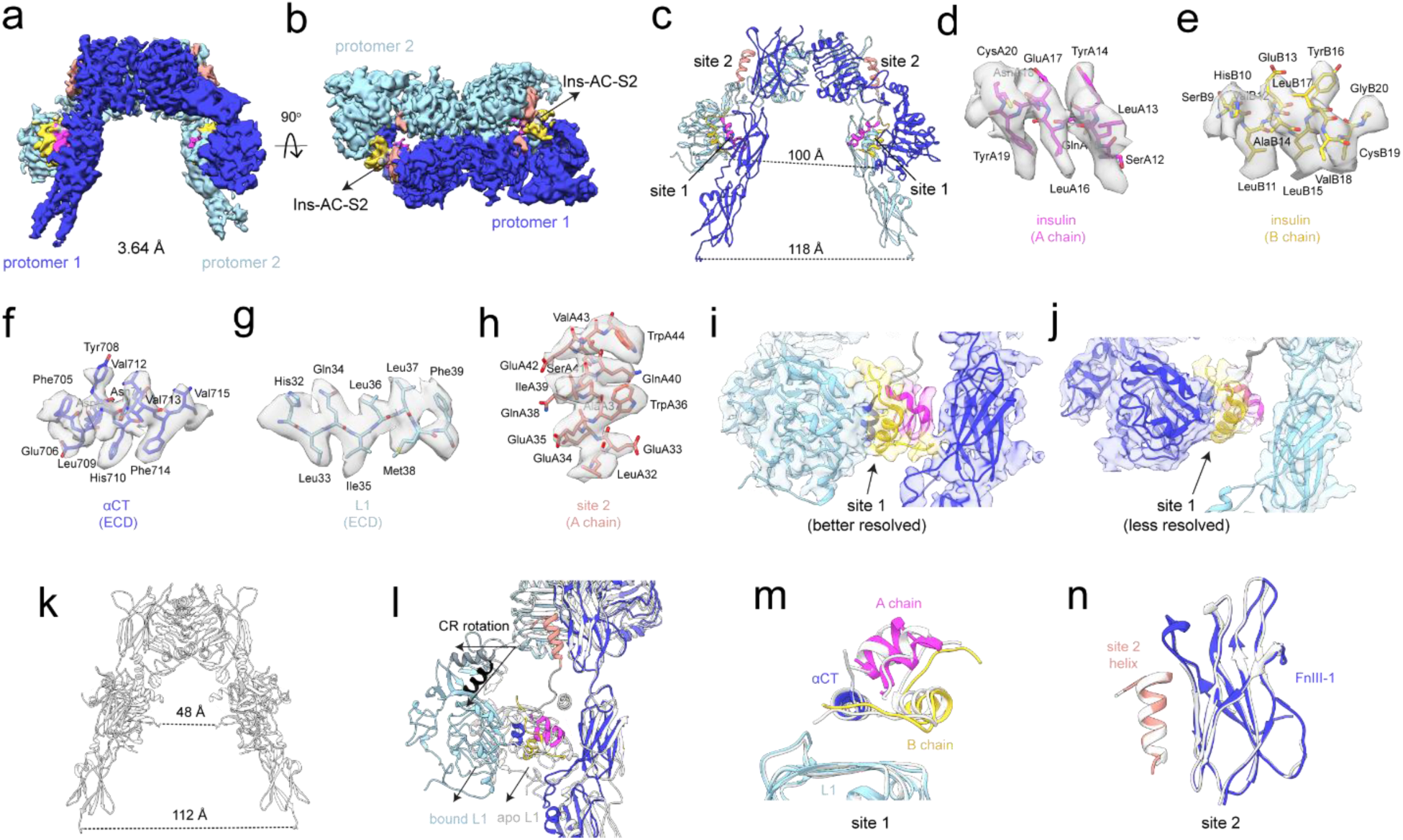
Structure of the ECD/Ins-AC-S2 complex. (a-b) Cryo-EM density map and model of ECD/Ins-AC-S2 determined at 3.64 Å from 314,550 particles showing (a) front view and (b) top view of the receptor. Protomers 1 and 2 are colored in dark and light blue, respectively. The density for Ins-AC-S2 is colored according to chain and binding segment (A chain insulin: magenta; A chain site 2 helix: salmon; B chain insulin: gold). (c) Model of the ECD/Ins-AC-S2 complex with distances shown between the fibronectin stalks (L909 to L909’) and L1 domains (E141 to E141’). (d-h) Representative density and model for site 1 (d) A chain insulin, (e) B chain insulin, (f) αCT, (g) L1, and (h) site 2-binding helix. (i-j) Model and map showing insulin from Ins-AC-S2 bound to site one on (i) the better reconstructed, and (j) the less well reconstruction receptor halves. (k) Model of the apo ECD (PDB: 4ZXB) with distances shown between the fibronectin stalks (L909 to L909’) and L1 domains (E141 to E141’). (l) Side view of IR/Ins-AC-S2 (color) overlayed with the apo ECD (white, PDB: 4ZXB) aligned to the FnIII-1 domains. The hinging rotation between CR and L2 domains in both models is shown with arrows. The first helices of the CR domains in the Ins-AC-S2 and apo models are colored in slate grey and black, respectively. (m) Alignment of ECD/Ins-AC-S2 with full-length IR from mouse complexed with insulin (white, PDB: 7STH) aligned on the L1 domains. (n) Alignment of ECD/Ins-AC-S2 with full-length IR from mouse complexed with the site 2 helix alone (white, PDB: 8DTM) aligned on the FnIII-1 domains.

Although less dramatic than for S961, the ECD/Ins-AC-S2 map also displayed asymmetry with one half of the receptor exhibiting reduced local resolution (Sup. Fig. 7). However, both fibronectin stalks were visualized, and the map was of sufficient resolution to create a model by rigid body fitting the extracellular domains of the receptor and Ins-AC-S2 (Fig. 5c). The resulting model clearly showed Ins-AC-S2 molecules bound to both sets of binding sites, resulting in a stoichiometry of 1:2 IR dimer:Ins-AC-S2. The more clearly reconstructed receptor half was of sufficient resolution to model many side chains for Ins-AC-S2 and ECD at the binding interfaces (Fig. 5d-h). Intriguingly, the overall conformation of the complex was somewhat asymmetric, with the insulins from Ins-AC-S2 molecules adopting slightly different positions at site 1 on either half of the receptor (Fig. 5i-j). At both sites, Ins-AC-S2 makes canonical contacts with L1 and αCT. However, on the better reconstructed receptor half, additional connecting density was observed between the insulin component of Ins-AC-S2 and the receptor FnIII-2 and insert (ID) domains. This new density was not observed on the less clearly reconstructed site 1 of the opposite receptor half, suggesting that interactions between the Ins-AC-S2 insulin and FnIII-2/ID may provide structural stability.

The ECD/Ins-AC-S2 complex adopts an overall conformation that deviates from other reported inverted-V conformations. Like the apo ECD structure, the fibronectin stalks are separated by 118 Å (Fig. 5c). However, the L1 and CR modules are raised away from the fibronectin stalks, creating an L1-L1 separation roughly twice that of the apo ECD structure (100 vs. 48 Å, Fig. 5c,k). Overlay of apo (4ZXB^7^) and Ins-AC-S2-bound ECD revealed that the wider L1-L1 separation results from hinging between the CR and L2 domains, positioning the L1 domains farther from the fibronectin stalks in the ECD/Ins-AC-S2 complex. This adjustment is required to accommodate the insulin of Ins-AC-S2 at site 1 (Fig. 5l).

Numerous published symmetric and asymmetric structures of the IR bound to 1-4 insulin molecules show engagement of L1 and αCT at site 1 and FnIII-1 at site 2^8–12,22–26^. To see if the Ins-AC-S2 insulin makes canonical contacts with IR site 1, we compared our structure to a published symmetric model of the FL-IR bound to two insulins^12^. As expected, Ins-AC-S2 at site 1 overlays well with a published IR/insulin structure when aligned to the L1 domain (Fig. 5m). We also compared our structure to the published structure of the isolated site 2 helix bound to FL-IR^19^. Again, we saw that the site 2 helix from Ins-AC-S2 overlays well with the published site 2 helix complexed structure (Fig. 5n). Thus, as for S961, the site 1 (insulin analogue) and site 2 (site 2 peptide) portions of Ins-AC-S2 each contact their binding surfaces on the receptor in the same manner as for active complexes, but their relative position in the sequence dictates an inactive inverted-V conformation for the receptor.

### A novel Ins-AC-S2 binding site stabilizes the antagonized receptor

Prompted by the conformational heterogeneity indicated by the cryo-EM map (Fig. 5i-j, Sup. Fig. 7), we explored dynamics of the ECD/Ins-AC-S2 complex using 3DVA. Five subsets of particles were obtained, and subsequent reconstructions revealed a subset of particles with strong density for the fibronectin stalk and a contrasting subset of particles with weaker density for the FnIII-3 domain of the fibronectin stalk (Fig. 6a). In the reconstruction with strong fibronectin stalk density, previously unreported contacts were apparent between the insulin of Ins-AC-S2 and the FnIII-2 and insert domain (ID) of the adjacent IR protomer. In contrast, these contacts were not apparent in the reconstruction with less density for the FnIII-3 domain. We call the new binding site between Ins-AC-S2 and IR site 1Ʌ, and we speculate it may stabilize the adjacent fibronectin stalk, contribute to binding, and hence reinforce the antagonistic activity of Ins-AC-S2.

**Figure 6:**
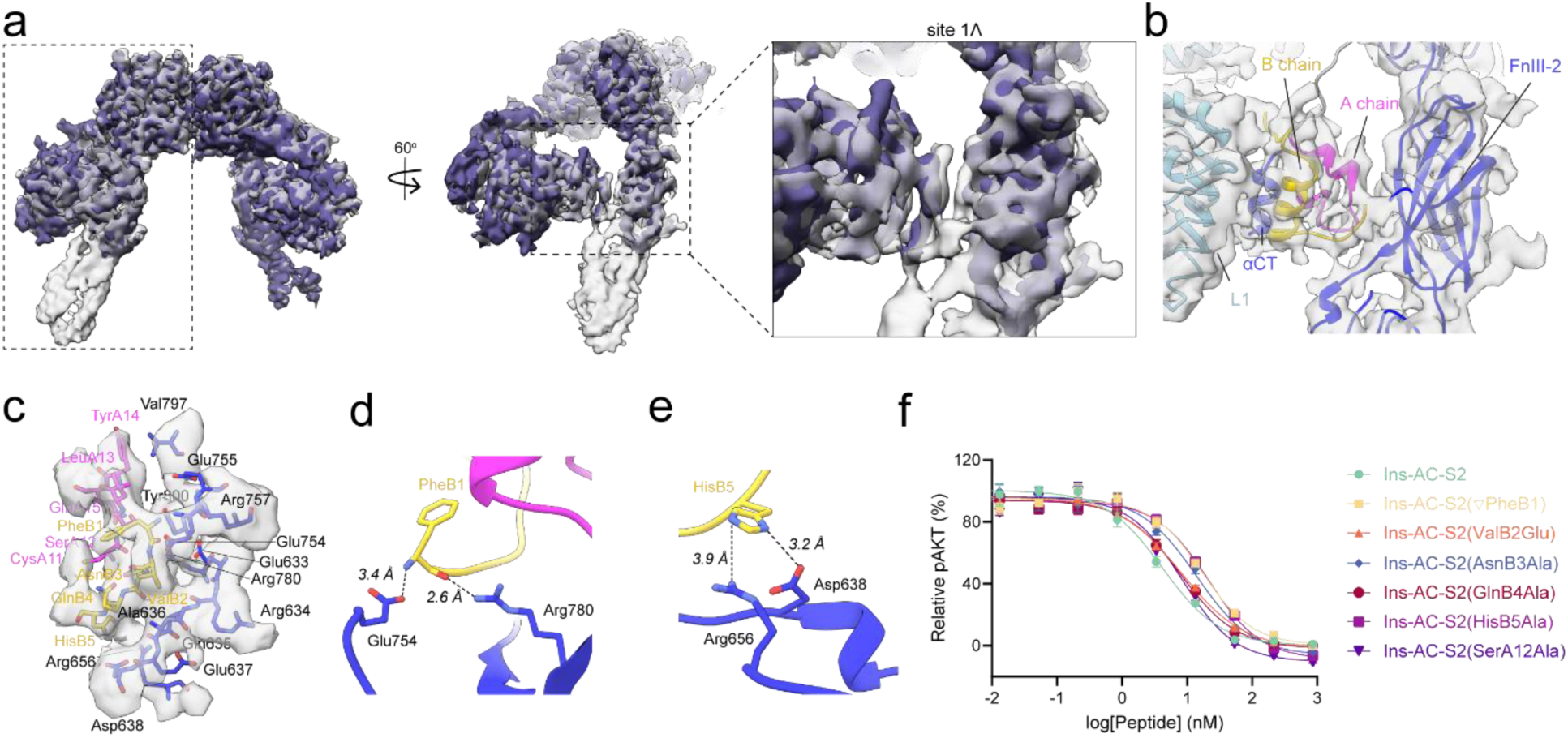
Structural and biochemical characterization of ECD/Ins-AC-S2 site 1Ʌ. (a) Overlay of the two most different density maps obtained from 3DVA with a focused mask on the receptor half containing site 1, site 1Ʌ, and site 2, showing front, side, and close up views of site 1Ʌ. Gray map at 3.83 Å (162,982 particles). Dark purple map at 3.85 Å (145,601 particles). (b) Map and model from local refinement of site 1Ʌ at 3.89 Å from 212,201 symmetry expanded particles. (c) Map and model of Site 1Ʌ showing all residues of Ins-AC-S2 and ECD within 5 Å of each other labeled. Ins-AC-S2 A chain, B chain, and ECD colored magenta, gold, and blue, respectively. (d-e) Site 1Ʌ side chain interaction pairs showing (d) a salt bridge between the N-terminal amide of PheB1 and ECD Glu754, and charge-polar interaction between ECD Arg780 and the PheB1 backbone carbonyl; and (e) hydrogen bonds between HisB5 and ECD Arg656 and Asp638. (f) *In vitro* pAKT assay evaluating the antagonistic potency of Ins-AC-S2 and its mutants against 43 nM human insulin in NIH 3T3 cells overexpressing human IR-B.

To better visualize site 1Ʌ, symmetry expansion, focused 3D classification, and local refinement were performed to obtain a 3.89 Å half map from a subset of 212,201 particles. Although this map had lower global resolution than the consensus map at 3.64 Å, it showed more detail and higher local resolution at site 1Ʌ (Fig. 6b, Sup. Fig. 8). This map allowed modeling of numerous sidechains at site 1Ʌ, including for the entire Ins-AC-S2 B chain N-terminus, which is rarely visible in IR/insulin structures due to its inherent flexibility but makes numerous contacts with FnIII-2/ID in our structure (Fig. 6c). Contacts include five residues in the Ins-AC-S2 A chain (residues 11-15), five residues at the B chain N-terminus (residues 1-5), and 13 residues from ECD FnIII-2 and ID (residues 633-638, 656, 754-755, 757, 780, 797, and 800). The site 1Ʌ interface is stabilized by numerous charge-charge, charge-polar, and hydrogen bonding interactions, most of which involve B chain residues (Fig. 6d-e).

To test the importance of site 1Ʌ in Ins-AC-S2 antagonism, we developed six Ins-AC-S2 mutants designed to weaken this interaction: ΔPheB1 (deletion of Phe1 from the B chain), ValB2Glu, AsnB3Ala, GlnB4Ala, HisB5Ala, and SerA12Ala. These mutants displayed ∼2-5-fold increase in IC_50_ values relative to WT Ins-AC-S2 (Fig. 6f, Table 1). The largest reduction in potency was observed for mutants ΔPheB1 and HisB5Ala, likely because these residues form the most direct interactions with the receptor at site 1Ʌ (Fig. 6d-e). Overall, the effects of mutating site 1Ʌ were modest, but since the contacts are only possible when the receptor is in the inactive inverted-V conformation and this interface supports antagonism, its stabilization may be an attractive route for developing future IR antagonists with increased potency.

**Table 1:** Antagonism of site 1Ʌ Ins-AC-S2 mutants. IC₅₀ values (nM) were determined by nonlinear regression analysis of dose–response curves using GraphPad Prism 9 (GraphPad Software, California, USA).

| Peptide | IC<sub>50</sub> ± SE (nM) |
| --- | --- |
| Ins-AC-S2 | 4.21 ± 0.63 |
| Ins-AC-S2($\nabla$PheB1) | 17.2 ± 2.2 |
| Ins-AC-S2(ValB2Glu) | 8.23 ± 0.93 |
| Ins-AC-S2(AsnB3Ala) | 14.0 ± 2.3 |
| Ins-AC-S2(GlnB4Ala) | 8.00 ± 1.02 |
| Ins-AC-S2(HisB5Ala) | 21.4 ± 2.1 |
| Ins-AC-S2(SerA12Ala) | 8.35 ± 1.21 |

## CONCLUSIONS

IR antagonists are a promising therapeutic approach for treatment of HI, and understanding the structural principles of IR antagonism is critical for development of next-generation designs. Here, we determined structures of the IR bound to two antagonists: S961 and Ins-AC-S2, both of which contain site 1-site 2 binding segment organization. Both antagonists bind IR in the inactive inverted-V conformation, explaining their antagonism. By comparing our S961/IR structure to the previously reported structure of mouse IR with S597^19^, a peptide that contains the same binding segments but in the opposite order, we determined that peptide binding site organization controls the position of the IR L1 domain relative to the adjacent FnIII-1 domain, which subsequently controls the overall conformation of the receptor. These findings provide an explanation for the relationship between peptide binding site organization and receptor activity.

Although both IR/antagonist complexes adopted inverted-V conformations, our structures revealed two striking differences between binding of S961 and Ins-AC-S2 to the receptor. First, the site 1 helix of S961 binds the exact same surface on L1 as αCT, thus creating competition between the ligand and the receptor itself, which likely has the undesirable effect of destabilizing the antagonized complex. Intriguingly, dissociation of S961 from site 1 was only observed on one half of the receptor. Since αCT bridges site 1 on each half of the receptor through disulfide bonds^27^, displacement of αCT upon binding of S961 at the first site 1 may change the affinity of S961 for the second site 1. This may explain the asymmetric conformational heterogeneity observed in the structure with potential relevance for the reported negative cooperativity of the receptor^28^. In contrast, Ins-AC-S2 engages αCT at site 1, reducing conformational heterogeneity and stabilizing the complex, which could explain its slightly higher potency. We expect that future antagonists designed to engage αCT will be most effective. The second critical difference between the antagonized complexes was the presence of a novel interaction at site 1 in the IR/Ins-AC-S2 complex. This site 1Ʌ interface can only exist when the receptor is in the inverted-V conformation and our structural and mutagenesis data indicate that it contributes to binding and stabilizing the antagonized receptor. Enhancing the site 1Ʌ interaction is a promising strategy for development of next-generation IR antagonists.

## METHODS

### Peptides

Purchased S961 acetate (MedChemExpress^®^) was solubilized using 10 mM NaOH, diluted 2-fold with buffer containing 20 mM HEPES, pH 7.5, 150 mM NaCl, 0.02% glyco-diosgenin (GDN) and stored at −80 °C. Concentrations were determined by absorbance at 280 nm using an extinction coefficient of 19,605 M^-1^.cm^-1^. Ins-AC-S2 was synthesized as described previously^15^. The powder was dissolved in 20 mM HEPES, pH 8.0, and stored at −80 °C. Concentrations were determined by absorbance at 280 nm using an extinction coefficient of 18,825 M^-1^.cm^-1^.

### Production of IR ECD

A polyclonal doxycycline-inducible adherent/suspension hybrid HEK293 stable cell line for production of IR-A ECD was produced using the PiggyBac transposase system^29^. Briefly, an IR transposition cassette flanked by piggyBac inverted repeat sequences containing a bidirectional tetracycline promotor, the gene for IR-A ectodomain (residues 1-930), a 7-residue S/G/P linker, a C-terminal His_6x_-tag followed by a 2x-FLAG tag, and a puromycin selection marker, was cloned into pTre-Bi mammalian expression vector using HiFi DNA assembly to produce the pTre-Bi-ECD plasmid. Adherent HEK293 cells were co-transfected with HyPBase-mCherry containing a transposase gene and pTre-Bi-ECD using ExpiFectamine™ 293, followed by several passages in the presence of 1 µg/mL. Expression was scaled up to 1 L in FS 293^TM^ media and induced with 2 µg/mL doxycycline. After 2 days, expression media containing secreted ECD was harvested and filtered using 0.45 µm filter paper. ECD was purified by Ni-NTA affinity chromatography and size-exclusion chromatography (SEC) using a Superose^TM^ 6 Increase 10/300 GL column pre-equilibrated with buffer containing 20 mM HEPES, pH 7.5, and 150 mM NaCl. Average protein purifications yielded ∼5.4 mg/L ECD that was flash frozen and stored at −80 °C.

### Production of full-length IR

A polyclonal doxycycline-inducible adherent/suspension hybrid HEK293 stable cell line for production of full-length IR-A was produced using the PiggyBac transposase system^29^ as described above. An IR transposition cassette flanked by piggyBac inverted repeat sequences containing a bidirectional tetracycline promotor, the gene for full-length IR-A with a C-terminal 2x-FLAG tag, and a puromycin selection marker, was cloned into pTre-Bi mammalian expression vector HiFi DNA assembly to produce the pTre-Bi-IR plasmid. Adherent HEK293 cells were co-transfected with HyPBase-mCherry containing a transposase gene and pTre-Bi-IR using ExpiFectamine™ 293. Following several rounds of puromycin selection and induction with 2 µg/mL doxycycline, cells expressing IR were labeled with Alexa Fluor 488-conjugated IR antibody and isolated using flow cytometry. Expression was scaled up to 1 L in Expi293^TM^ suspension media supplemented with 10 mM sodium butyrate to promote expression and induced with 2 µg/mL doxycycline for 2 days. Harvested cells were lysed by sonication in buffer containing 20 mM HEPES, pH 7.5, 400 mM NaCl, 5% glycerol, 1% n-dodecyl-beta-maltoside (DDM), and 0.1% cholesteryl hemisuccinate (CHS), followed by stirring at 4 °C for 3 hours to allow for membrane protein extraction. Membrane extracted IR was purified from cleared cell lysate using FLAG affinity chromatography and eluted with 3x-FLAG peptide. Due to incomplete cellular receptor processing, FLAG affinity was followed by incubation with 5 units/mL furin protease (NEB^®^) for 5 hours at room temperature, followed by SEC using a Superose^TM^ 6 Increase 10/300 GL column pre-equilibrated with buffer containing 20 mM HEPES, pH 7.5, 150 mM NaCl, supplemented with 0.03% DDM and 0.003% CHS or 0.02% GDN. Average protein purifications yielded ∼0.6 mg/L IR that was flash frozen and stored at −80°C.

### Cryo-EM

#### IR/S961

The IR/S961 map was obtained from several protein preparations and datasets. SEC-purified FL-IR at 13.5 µM or ECD at 14.7 µM supplemented with 4 mM CHAPSO were incubated for 30-60 min with 29.6 µM or 22 µM S961, respectively. AuFoil, Quantifoil R1.2/1.3, 300 mesh grids, or nanowire grids were glow discharged for 25 s or 2 min on one side, or 2 min on each side, respectively. Two to 3.5 uL samples were applied to AuFoil and Quantifoil grids, blotted for 3-7 s at 11 °C with 80% relative humidity and flash frozen in liquid ethane using a Leica EM GP2 automatic plunge freeze instrument. One microliter sample was applied to nanowire grids and flash frozen using our homemade ultrafast grid plunger^30^. The piezo atomizer was energized for 20 ms prior to plunging, resulting in a total application-to-freeze time of 250 ms as determined by high frame rate video. Data were collected on a ThermoFisher Titan Krios G3 transmission electron microscope equipped with a Gatan BioQuantum K3 energy filter and direct electron detector at 0 or 15° tilt angles. Movies were collected at 81,000x magnification with a pixel size of 1.06 Å/pixel and an electron dose of 50 e^-^/A^2^ over 40 frames/exposure with a defocus range of −0.8 to −2.0 µm using EPU software.

Data processing was performed using CryoSPARC v4.7^31^ (Sup. Fig. 1). 21,434 movies were combined from datasets collected from five different grids. Patch motion correction, CTF estimation, template picker, and particle extraction using a box size of 320 px down sampled to 256 px for initial processing were performed on each dataset individually. The resulting 7,329,122 particles were subjected to iterative rounds of 2D classification, followed by heterogenous refinement into six classes. Non-uniform refinement using C2 symmetry with marginalization was performed using the best 745,509 particles and produced a 3.78 Å reconstruction. This particle set was then subjected to two separate data processing workflows. First, particles were re-extracted using a box size of 320 px, followed by heterogeneous refinement into five classes. The 378,182 particles from the best class produced a final reconstruction of 3.68 Å from non-uniform refinement using C2 symmetry with marginalization. A model was built into this map starting from PDBs 7SL1^12^, 5J3H^17^, and 8DTM^19^ with minor adjustments to fit the density using UCSF Chimera^32^ and ISOLDE^33^ in UCSF ChimeraX^34^. Refinements were performed using Phenix^35,36^ and validated using MolProbity^37,38^ (Sup. Table 1). Local refinement with focused masks on each half of the receptor were performed to assess the local resolution of each receptor half while eliminating bias introduced by global particle alignment. In addition, 3DVA^39^ was performed on the larger particle set (745,509 particles). The two most different subsets of particles obtained from 3DVA were further refined using non-uniform refinement using C2 symmetry with marginalization, resulting in 3.99 Å and 3.97 Å resolution reconstructions. Resolutions were determined using an FSC cutoff of 0.143.

#### FL-IR/Ins-AC-S2

Purified FL-IR was prepared in 20 mM HEPES, pH 7.5, 150 mM NaCl and solubilized by 0.02% GDN, or 0.03% DDM and 0.003% CHS. For some samples, detergent was replaced with the amphipathic membrane scaffold protein, MSP1E3D1 using previously published protocols for nanodisc assembly^40,41^, although lipid disc density was not observed in the cryo-EM data (data not shown). FL-IR samples solubilized by MSP1E3D1 were prepared at 1 µM with 2.2 µM Ins-AC-S2. FL-IR samples solubilized by detergent were prepared at 5.3 µM, 13 µM, 14 µM, or 16 µM and incubated with 11.7 µM, 26 µM, 28 µM, or 35.2 µM Ins-AC-S2, respectively for 30-60 min. Quantifoil R1.2/1.3, 300 or 400 mesh grids were glow discharged for 7 or 25 s. 3.5 uL samples were applied to grids, blotted for 5.5-7 s at 11 °C with 80% relative humidity and flash frozen in liquid ethane using a Leica EM GP2 automatic plunge freeze instrument. Nanowire grids prepared on our homemade ultrafast grid plunger^30^ were glow discharged for 2 min on each side, and 1 or 1.5 µL sample was applied. The piezo atomizer was energized for 20 ms prior to the start of plunging, resulting in a total application-to-freeze time of 250 ms as determined by high frame rate video. Data were collected on a ThermoFisher Titan Krios G3 transmission electron microscope equipped with a Gatan BioQuantum K3 energy filter and direct electron detector. Movies were collected at 81,000x magnification with a pixel size of 1.06 Å/pixel and an electron dose of 50 e^-^/A^2^ over 40 frames/exposure with a defocus range of −0.8 to −2.0 µm using EPU software.

Data processing was performed using CryoSPARC v4.7^31^ (Sup. Fig. 2). 14,042 movies were combined from datasets collected on six different grids. Patch motion correction and CTF estimation were performed on each dataset individually. Particles from one dataset of 768 movies were picked using blob picker with minimum and maximum particle diameters of 160 and 200 Å, respectively. Particle extraction with a box size of 320 px was followed by three rounds of 2D classification. The best 2D classes (40,898 particles) were used as templates for particle picking from the combined micrographs. Template picked particles were extracted using a box size of 360 px, followed by 10 rounds of 2D classification, particle re-extraction, a final round of 2D classification, and non-uniform refinement with C1 symmetry of the best 475,634 particles, resulting in a 6.44 Å reconstruction. Extensive downstream data processing (3D classification, heterogenous refinement, and 3DVA^39^) was performed in attempt to obtain a higher resolution map but without success.

#### ECD/Ins-AC-S2

Purified ECD at concentrations of 1.5 µM, 12.6 µM, or 14.8 µM was incubated with Ins-AC-S2 at concentrations of 3.3 µM, 50 µM, or 58 µM, respectively, in buffer composed of 20 mM HEPES, pH 7.5, and 150 mM NaCl, with 0 mM, 4 mM or 8 mM CHAPSO. Quantifoil R1.2/1.3 or R2/2, 300 mesh grids were glow discharged for 25 s or 2 min. After complex incubation for 30-60 min, 3.5 uL samples were applied to grids, blotted for 6-7.5 s at 11 °C with 80% relative humidity and flash frozen in liquid ethane using a Leica EM GP2 automatic plunge freeze instrument. Data were collected on a ThermoFisher Titan Krios G3 transmission electron microscope equipped with a Gatan BioQuantum K3 energy filter and direct electron detector at 0 or 30° tilt angles. Movies were collected at 81,000x magnification with a pixel size of 1.06 Å/pixel and an electron dose of 50 e^-^/A^2^ over 40 frames/exposure with a defocus range of −0.8 to −2.0 µm using EPU software.

Data processing was performed using CryoSPARC v4.7^31^. 24,583 movies were combined from datasets collected on five different grids (Sup. Fig. 3). Patch motion correction and CTF estimation were performed on each dataset individually, followed by template particle picking using templates generated from the FL-IR/Ins-AC-S2 volume. Particles were extracted using a box size of 320 px down sampled to 256 px for initial processing. Following iterative rounds of 2D and 3D classification, the 1,645,091 best particles from the combined datasets were pooled and subjected to one final round of 2D classification. The best classes (1,148,473 particles) were selected and used to generate a 3.90 Å reconstruction using non-uniform refinement using C2 symmetry with marginalization. Additional junk particles were removed through heterogeneous refinement with six classes, followed by particle re-extraction with a box size of 320 px, reference-based motion correction, and a final round of heterogeneous refinement with six classes. Non-uniform refinement of the best 314,550 particles resulted in a 3.64 Å final reconstruction.

To characterize the dynamics of the complex, two rounds of 3DVA were performed with a filter resolution of 10 Å and focused masks around each half of the receptor divided along its C2 symmetry axis. The two most different subsets of particles were further refined using non-uniform refinement using C2 symmetry with marginalization, resulting in 3.83 Å and 3.85 Å reconstructions from 162,982 particles and 145,601 particles, respectively. To obtain a high-resolution map of the Ins-AC-S2 insulin site 1Ʌ interaction, symmetry expansion was performed on the 314,550 particles used in the final reconstruction of the full complex, followed by focused 3D classification and local refinement, yielding a half map with a resolution of 3.89 Å from 212,201 symmetry expanded particles. Resolutions were determined using an FSC cutoff of 0.143. Model building was performed starting from PDBs 7MD4^42^, 6PXV^10^, and 8DTM^19^ with minor adjustments to fit the density using UCSF Chimera^32^ and ISOLDE^33^ in UCSF ChimeraX^34^ (Sup. Table 1). Refinements were performed using Phenix^35,36^ and validated using MolProbity^37,38^.

### Cell-Based pAKT Assays

Antagonistic activities of Ins-AC-S2 and its mutants were evaluated using a cell-based pAKT (Ser473) assay. Endogenous levels of pAKT were measured in a human IR-B overexpressing NIH 3T3 cell line, derived from IGF-1R knockout mice (a generous gift from A. Morrione, Thomas Jefferson University). Cells were cultured in DMEM (Gibco) with 10% fetal bovine serum (FBS, Gibco), 100 U/mL penicillin–streptomycin (Gibco), 2 μg/mL puromycin (Gibco), and 0.1 mg/mL normocin (InvivoGen) at 37 °C under 5% CO_2_. For the assays, 30,000 cells per well and 100 μL per well, were plated in a 96-well plate with culture media containing 1% FBS. 20 h later, the media was replaced with 50 μL of culture media containing different concentrations of Ins-AC-S2 or its mutants in the presence of 43 nM human insulin. After treatment at 37 °C for 25 min, the solution was removed and the HTRF pAKT Ser473 kit (Cisbio) was used to measure the phosphorylation of AKT. Briefly, the cells were first treated with cell lysis buffer (50 μL per well). After mild shaking for 30-45 min, 16 μL of the cell lysate was added to 4 μL of the detecting reagent in a white 384-well plate. After 4 h of incubation, the plate was read in a SpectraMax iD5 plate reader (Molecular Devices) and the data were processed according to the manufacturer’s protocol. All data points are shown as mean ± SEM (n = 4).

## DATA AVAILABILITY

The cryo-EM map and model for the IR/S961 complex has been deposited into the EMDB/PDB database under the entry ID: PDB ID 9PUV, Extended PDB ID pdb_00009PUV, EMD-71877. The cryo-EM map and model for the ECD/Ins-AC-S2 complex has been deposited into the EMDB/PDB database under the entry ID: PDB ID 9PUW, Extended PDB ID pdb_00009PUW, EMD-71878. The cryo-EM half map and model for the ECD/Ins-AC-S2 complex focused on site 1Ʌ has been deposited to the EMDB/PDB database under the entry ID: PDB ID 9PVO, Extended PDB ID pdb_00009PVO, EMD-71894.

## ACKNOWLEDGMENTS

This research was funded by NIH/NIDDK R01DK127268 and the P&F grant from the Stanford Diabetes Research Center (P30DK116074). We thank Stanford Maternal & Children Research Institute support for Y. D. (Postdoc Fellowship) and D. H. C. (Pilot grant). We thank Breakthrough T1D for support for A.V. (Postdoc Fellowship). We thank David Belnap, Barbie Ganser-Pornillos, and the University of Utah Arnold and Mabel Beckman Center for Cryo-EM data collection and support. Cryo-EM data processing was made possible by resources from the Center for High Performance Computing at the University of Utah. We thank the University of Utah HSC Flow Cytometry Core Facility for cell sorting experiments. We thank Heidi Schubert (University of Utah) for helpful conversations.

## Contributions

A.V. and A.B. designed the study, prepared cryo-EM samples, collected cryo-EM data, and performed cryo-EM data processing and analysis. Y.D. prepared Ins-AC-S2 ligands, performed cell activity assays, and analyzed IR antagonism data. A.V. wrote the manuscript and prepared the figures with input and editing from all authors. D.C. and C.P.H supervised experimental design, data interpretation, and provided project funding. All authors provided intellectual insight.

## MATERIALS AND CORRESPONDENCE

Correspondence addressed to Christopher P. Hill.

