## Supplementary Information for "Structural basis of insulin receptor antagonism by bivalent site 1-site 2 ligands"

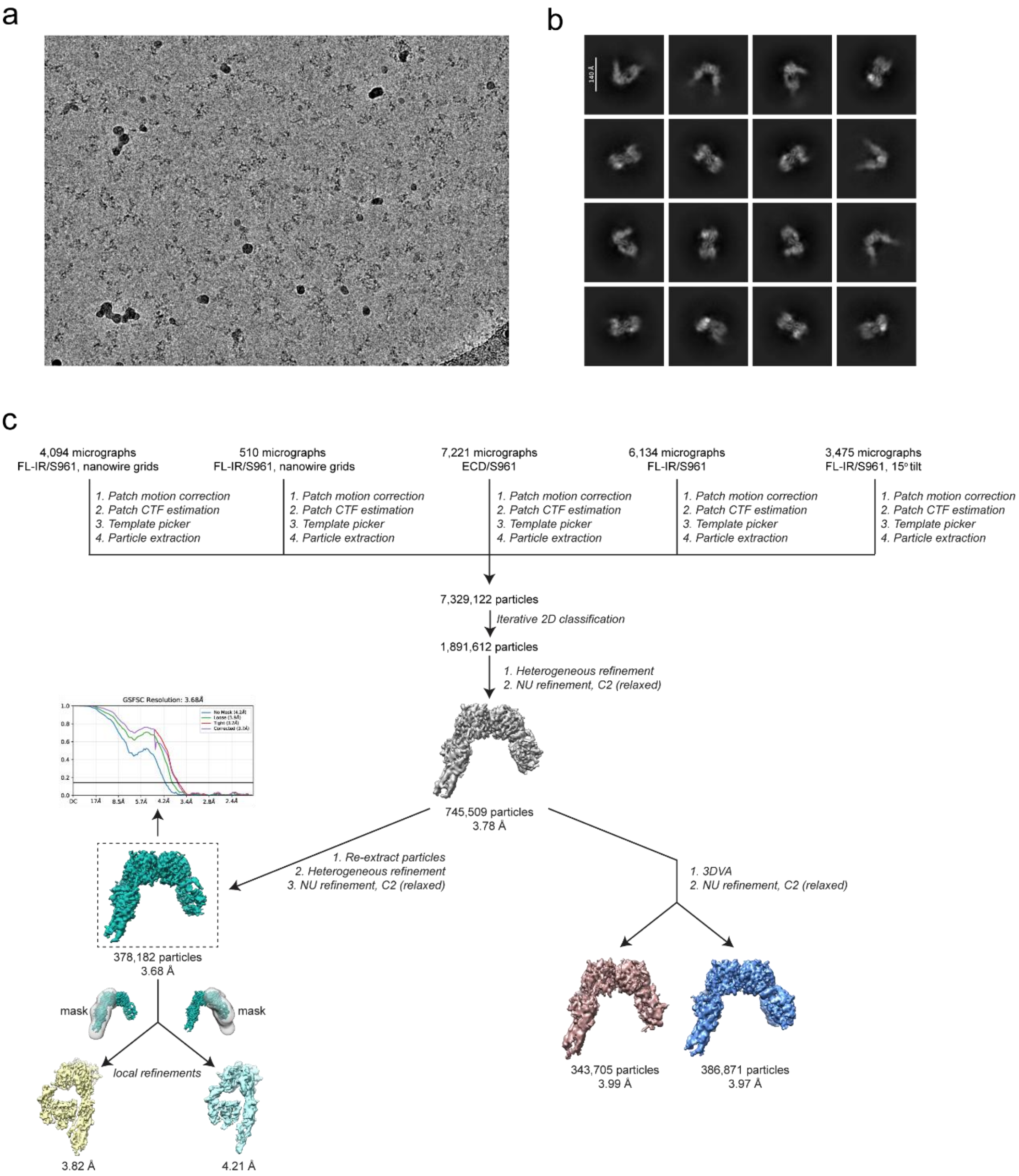


Sup. Fig. 1: IR/S961 data processing workflow. (a) Representative micrograph from the IR/S961 datasets. (b) Sixteen representative 2D class averages from the final subset of particles used in the IR/S961 reconstruction. (c) Data processing workflow performed in cryoSPARC to obtain a final reconstruction at 3.68 Å resolution from 378,182 particles. Particles from five datasets were used. The FSC plot for the consensus refinement used for molecular modeling is shown.

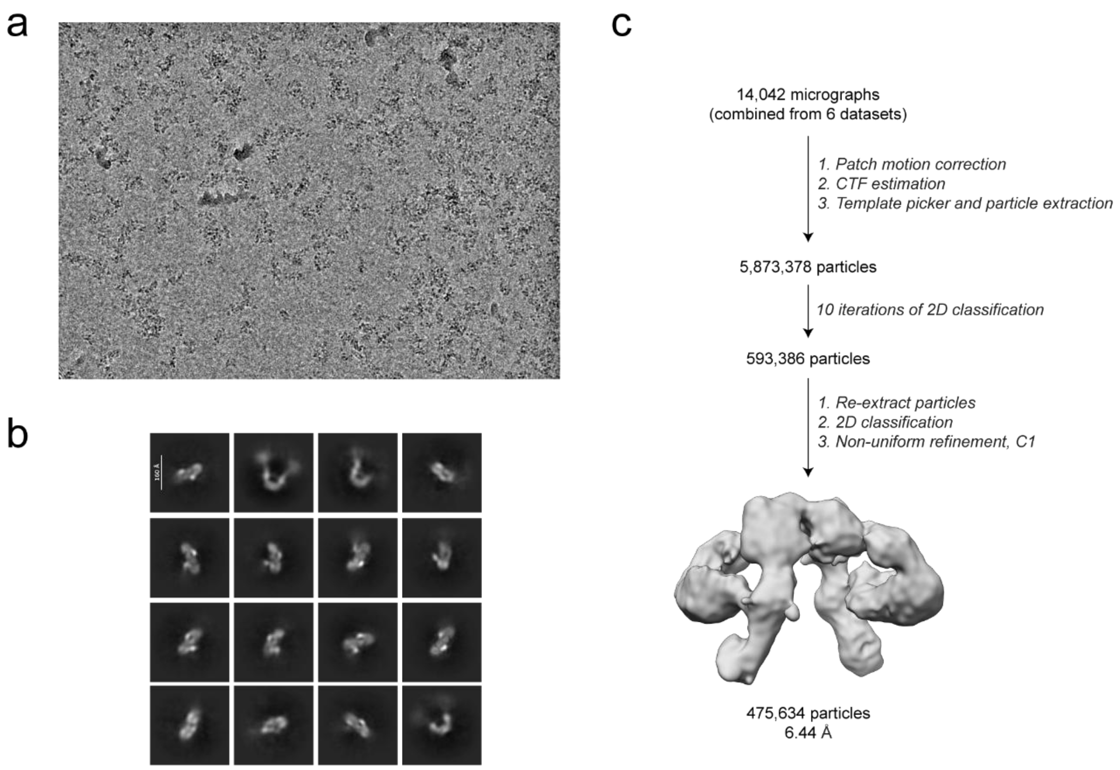


Sup. Fig. 2: FL-IR/Ins-AC-S2 cryo-EM data processing workflow. (a) Representative micrograph from the FL-IR/Ins-AC-S2 datasets. (b) Sixteen representative 2D class averages from the final subset of particles used in the FL-IR/Ins-AC-S2 reconstruction. (c) Data processing workflow performed in cryoSPARC to obtain a final reconstruction at 6.44 Å resolution from 475,634 particles.

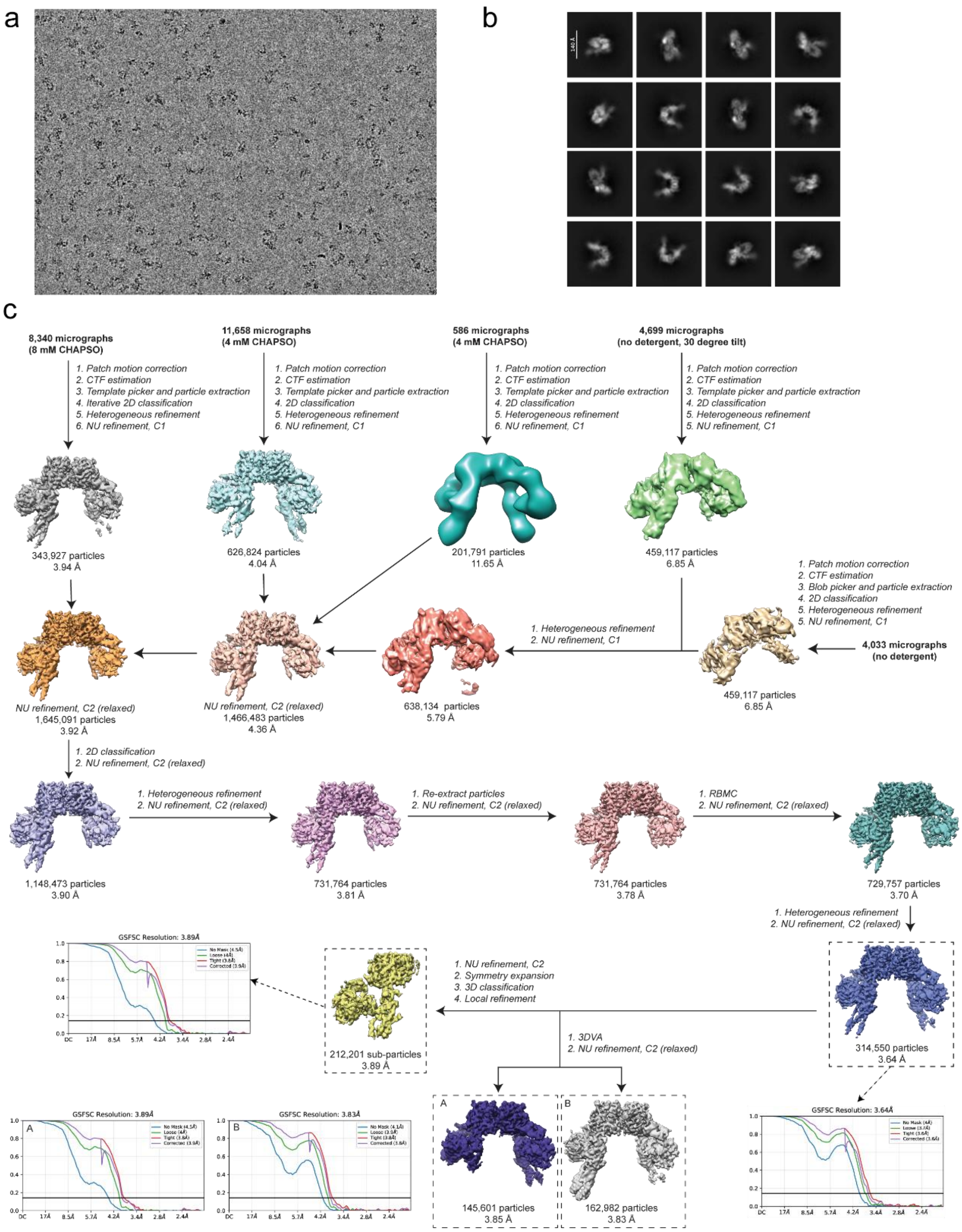


Sup. Fig. 3: Cryo-EM data processing workflow for the ECD/Ins-AC-S2 complex. (a) Representative micrograph from the ECD/Ins-AC-S2 datasets. (b) Sixteen representative 2D class averages from the final subset of particles used in the ECD/Ins-AC-S2 reconstruction. (c) Data processing workflow performed in cryoSPARC to obtain a final reconstruction at 3.64 Å resolution from 314,550 particles. Particles from five datasets were used. Maps used in final reconstructions and for building models are boxed, and the corresponding FSC curves are shown.

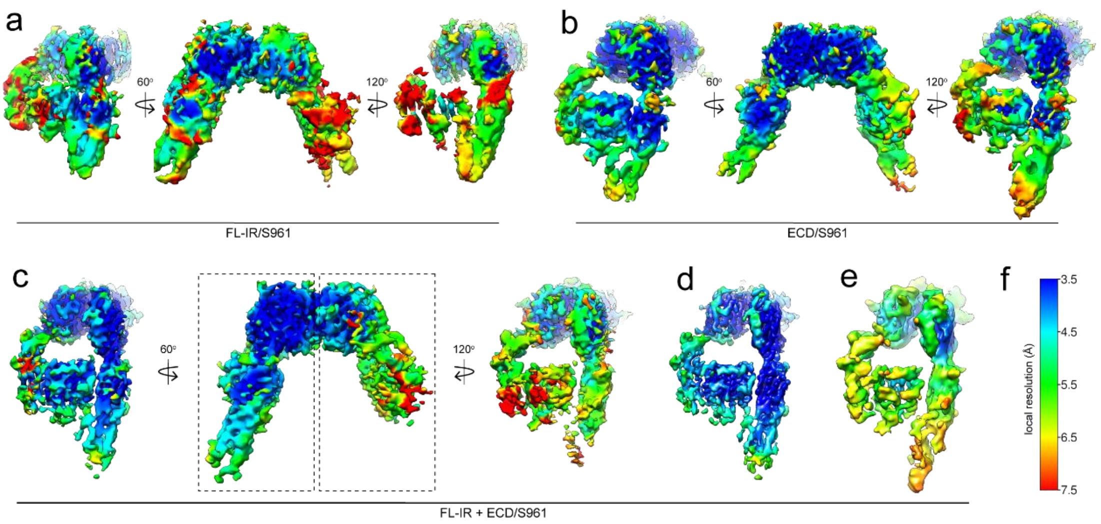


Sup. Fig. 4: IR/S961 maps colored by local resolution. (a) Three views of the best map obtained of the FL-IR/S961 complex determined to 3.88 Å resolution from 497,687 particles. (b) Three views of the best density map obtained of the ECD/S961 complex determined to 3.93 Å map from 248,634 particles. (c) Three views of the IR/S961 consensus map at 3.68 Å resolution from 378,182 particles from combined FL-IR/S961 and ECD/S961 datasets. (d-e) Side views of local refinements performed on the combined FL-IR/S961 and ECD/S961 datasets with focused masks on (d) the better resolved receptor half (3.82 Å), and (e) the less resolved receptor half (4.21 Å). (f) Local resolution color scheme used to color panels (a-e).

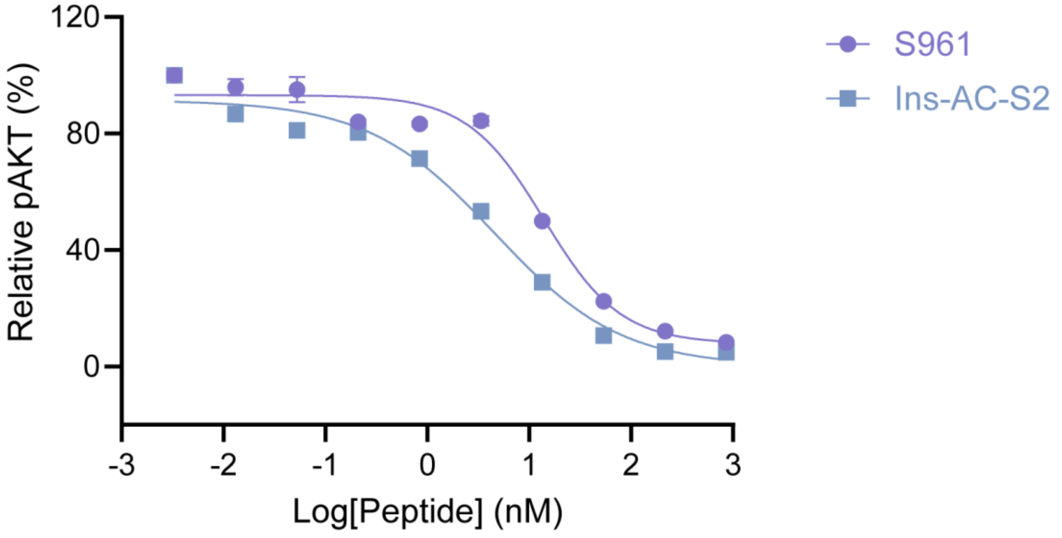


Sup. Fig. 5: Comparison of S961 and Ins-AC-S2 antagonism. *In vitro* pAKT assay evaluating the antagonistic potency of Ins-AC-S2 and S961 against 43 nM human insulin in NIH 3T3 cells overexpressing human IR-B.

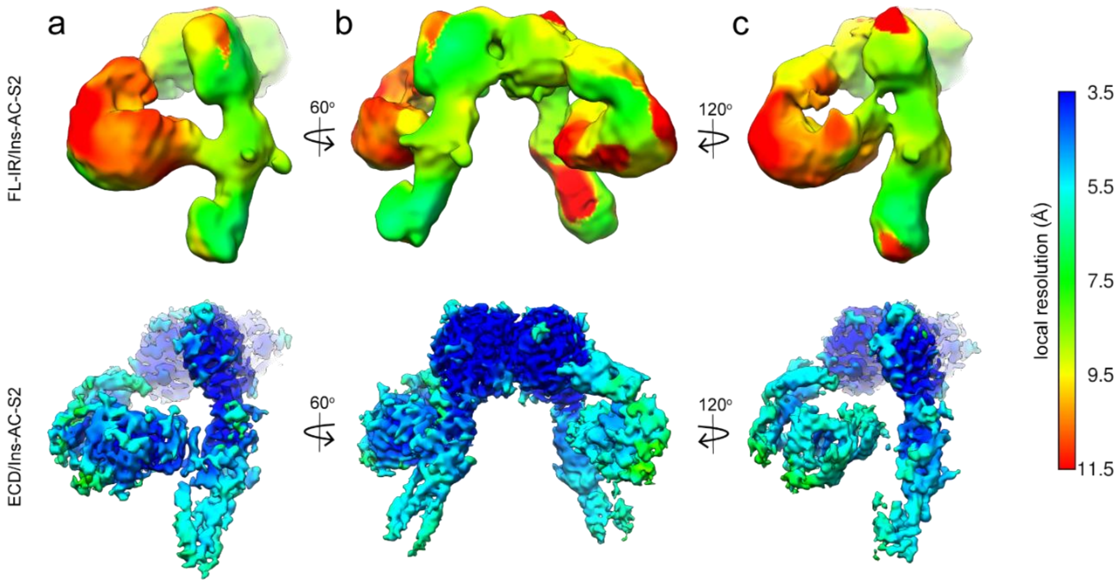


Sup. Fig. 6: Comparison of the FL-IR/Ins-AC-S2 and ECD/Ins-AC-S2 complexes. Three views of best density maps colored by local resolution of FL-IR (top) and ECD (bottom) bound to Ins-AC-S2 determined at 6.44 Å from 475,634 particles and 3.64 Å from 314,550 particles, respectively.

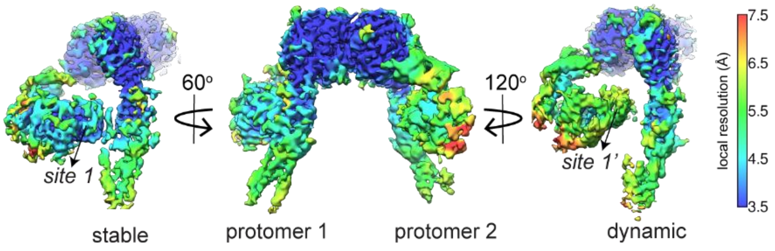


Sup. Fig. 7: Local resolution of the consensus ECD/Ins-AC-S2 density map. Cryo-EM density map of the ECD/Ins-AC-S2 complex determined to 3.64 Å colored by local resolution showing three views.

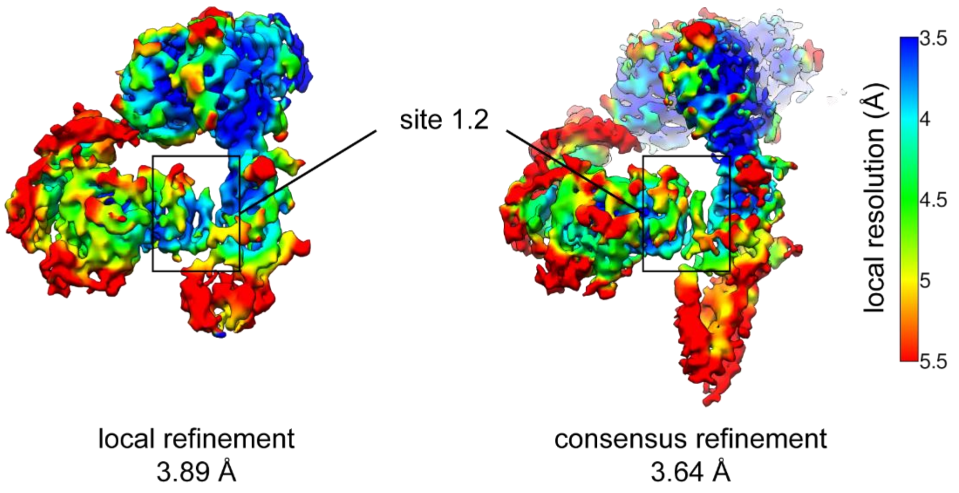


<u>Sup. Fig. 8: Local resolution of ECD/Ins-AC-S2 maps focused on site 1Λ.</u> Local refinement following symmetry expansion and 3D classification (left) and highest resolution non-uniform refinement (right). Density for site 1Λ is boxed.

Sup. Table 1: Cryo-EM data refinement statistics.

| | IR/S961 | ECD/Ins-AC-S2 (full model) | ECD/Ins-AC-S2 (site 1Λ) |
|---|---|---|---|
| EM Databank Accession ID | EMD-71877 | EMD-71878 | EMD-71894 |
| PDB Accession ID | 9PUV | 9PUW | 9PVO |
| **Data Collection/Processing** | | | |
| Nominal Magnification | 81,000x | 81,000x | 81,000x |
| Defocus Range (µm) | -0.8 to -2.0 µm | -0.8 to -2.0 µm | -0.8 to -2.0 µm |
| Number of micrographs | 21,434 | 24,583 | 24,583 |
| Initial particles | 7,329,122 | 7,042,324 | 7,042,324 |
| Symmetry imposed | C2 (relaxed) | C2 (relaxed) | C1 |
| Final Particles | 378,182 | 314,550 | 212,201 (symmetry expanded) |
| Map Resolution FSC 0.143 | 3.68 Å | 3.64 Å | 3.89 Å |
| **Validation** | | | |
| $CC_{box}$, $CC_{mask}$, $CC_{volume}$ | 0.75, 0.71, 0.70 | 0.72, 0.71, 0.70 | 0.72, 0.75, 0.74 |
| Atoms (non-H) | 26,242 | 26,929 | 12,699 |
| Protein residues | 1,654 | 1,697 | 801 |
| **Bonds (RMSD)** | | | |
| Length (Å) | 0.003 | 0.003 | 0.004 |
| Angles (°) | 0.529 | 0.538 | 0.591 |
| MolProbity score | 2.03 | 2.06 | 2.00 |
| Clash score | 10.1 | 9.51 | 8.43 |
| **Ramachandran plot (%)** | | | |
| Outliers | 0.06 | 0.18 | 0.13 |
| Allowed | 8.59 | 10.09 | 9.52 |
| Favored | 91.35 | 89.73 | 90.35 |
| Rotamer outliers (%) | 0.34 | 0.26 | 0.14 |
| CaBLAM outliers (%) | 3.8 | 4.23 | 6.86 |

Sup. Table 2: Side-by-side antagonism of S961 and Ins-AC-S2. $IC_{50}$ values (nM) were determined by nonlinear regression analysis of dose–response curves using GraphPad Prism 9 (GraphPad Software, California, USA).

| Peptide | $IC_{50}$ ± SE (nM) |
|---|---|
| S961 | 14.0 ± 1.7 |
| Ins-AC-S2 | 4.60 ± 0.70 |

#### Chemical Synthesis of Ins-AC-S2 and Its Mutants

*I. Reagents and Materials*

All commercially available chemical reagents, including those from Fisher, ChemPep, PurePep, CHEM-IMPEX, Novabiochem, Oakwood Chemical, Sigma Aldrich, Acros, TCI, and Adamas, were used without further purification. Solvents used in the experiments were either HPLC or reagent grade, sourced from Fisher Chemical and Sigma Aldrich. Ultrapure deionized water was obtained using the Milli-Q IQ 7000 water purification system (Merck, Darmstadt, Germany). 2-Chlorotrityl chloride (2-CTC) resin and Rink amide resin were purchased from ChemPep. Native human insulin was purchased from Invitrogen Life Technologies (Carlsbad, CA, USA). Before HPLC purification, crude peptides were pre-filtered using a Basix™ 13 mm syringe filter with a 0.45 μm pore size.

*II. High-Performance Liquid Chromatography (HPLC)*

HPLC-MS analysis was conducted on Agilent 6120 Quadrupole LC/MS and Agilent G6160A Quadrupole LC/MS, utilizing a Phenomenex C18 column (Jupiter® 50 × 2 mm, 5 μm, 300 Å) or Phenomenex C4 column (Jupiter® 50 × 2 mm, 5 μm, 300 Å) at a flow rate of 0.4 mL/min. The mobile phase comprised 0.1% (*v/v*) formic acid in water (solvent A) and 0.1% (*v/v)* formic acid in acetonitrile (solvent B). UV detection was performed at wavelengths of 220 nm, 240 nm, 260 nm and 280 nm, with the column temperature maintained at 40 °C.

Purification of all crude peptides and proteins was carried out using an Agilent 1260 HPLC system with a mobile phase consisting of 0.1% (*v/v*) TFA in water (solvent A) and 0.1% (*v/v*) TFA in acetonitrile (solvent B). The system was equipped with Agilent 1260 Infinity Quaternary pumps and a 1260 Infinity II UV detector, with detection wavelengths set at 220 nm, 240 nm, 260 nm and 280 nm. Separation was conducted using a Phenomenex C18 column (Luna® 250 × 10 mm, 5 μm, 300 Å) with a flow rate of 4 mL/min.

*III. General Procedures of Peptide Synthesis*

Peptide synthesis was performed using Fmoc-based solid-phase peptide synthesis (SPPS) on a Multiple Synthesizer SYRO I (MultiSynTech GmbH, Witten, Germany) equipped with a vortex stirring system. The synthesis followed a standard protocol: Fmoc groups were removed by treating the resin twice with 20% 4-methylpiperidine in DMF (1.2 mL) for 10 minutes at room temperature. After deprotection, the resin was thoroughly washed with DMF (1.3 mL × 5). Peptide elongation was achieved by coupling Fmoc-protected amino acids (5 equiv.) with HATU (4.8 equiv.) and DIPEA

(10 equiv.) in DMF (2.5 mL). Coupling reactions were conducted at 50 °C for cysteine and histidine, while all other amino acids were coupled at 70 °C for 10 minutes. Between each coupling and deprotection cycle, the resin underwent multiple DMF wash steps (1.3 mL × 3) to ensure efficient removal of unreacted reagents.

Before cleavage, the resin was thoroughly washed with DCM (5 mL × 3) and then placed under vacuum for one hour to ensure complete drying. Peptide cleavage and global deprotection were performed using TFA-based cleavage cocktails (TFA:TIPS:$H_2O$ = 95:2.5:2.5, *v/v/v*) for 2 hours. Following cleavage, the peptide-containing solution was separated from the resin by filtration, and the peptides were precipitated by adding the solution dropwise into cold diethyl ether (40 mL). The precipitate was collected by centrifugation at 3,500 × g for 3 minutes, after which the supernatant was discarded. The resulting pellet was further washed with diethyl ether (40 mL × 2) to remove residual impurities. The crude peptides were then dried under vacuum for at least 30 minutes, dissolved in a solution of MeCN/$H_2O$ containing 5% AcOH, analyzed via LC-MS, and purified using preparative HPLC.

*IV. Recombinant Expression of Sortase A*

Sortase A containing five amino acid mutations P94R, D160N, D165A, K190E and K196T was expressed in *E. coli* following David R. Liu's protocol[1]. E. coli BL21(DE3) transformed with pET29 Sortase A expression plasmids were cultured at 37 °C in LB with 50 µg/mL kanamycin until OD600 = 0.5-0.8. IPTG was added to a final concentration of 0.4 mM and protein expression was induced for three hours at 30 °C. Cells were harvested by centrifugation and resuspended in lysis buffer (50 mM Tris pH 8.0, 300 mM NaCl supplemented with 1 mM $MgCl_2$, 2 units/mL DNAseI (NEB), 260 nM aprotinin, 1.2 µM leupeptin, and 1 mM PMSF). Cells were lysed by sonication and the clarified supernatant was purified on Ni-NTA agarose following the manufacturer's instructions. Fractions that were >95% purity, as judged by SDS-PAGE, were consolidated and dialyzed against Tris-buffered saline (25 mM Tris pH 7.5, 150 mM NaCl). The concentration of Sortase A was determined by Nanodrop (extinction coefficient of 17,420 $M^{-1}$ $cm^{-1}$) and mixed with 20% glycerol, then stored at -80 °C freezer.

*V. Sortase A-Mediated Ligation of Ins-AC mutants and site 2 peptide*

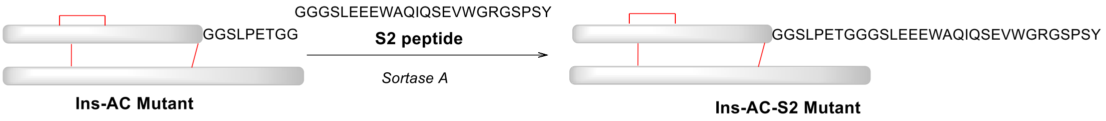


Ins-AC Mutant and S2 peptide were dissolved independently in reaction buffer (50 mM Tris·HCl, 150 mM NaCl, 5 mM CaCl2, pH 7.5) to a concentration of 1 mM. Then 1 equiv. of Ins-AC Mutant (100 μM) and 2 equiv. of S2 peptide (200 μM) were mixed in a 15 mL Eppendorf tube, 0.05 equiv. of Sortase A (5 μM) was added and reacted for 5 minutes, the reaction was quenched by a solution of MeCN/H2O (+ 5% AcOH), and analyzed by LC-MS. The resulting mixture was then purified using semi-preparative HPLC (Phenomenex C18 column, Luna® 250 × 10 mm, 5 μm, 300 Å, linear gradient 28% to 40% solvent B over 24 min), the fractions of desired Ins-AC-S2 mutant were collected and lyophilized to yield the product as a white powder.

### Characterization of Ins-AC-S2 and Its Mutants

***<u>Ins-AC( ∇PheB1)</u>***

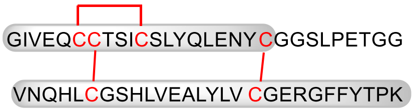


Chemical Formula: $C_{271}H_{411}N_{71}O_{84}S_6$
Exact Mass: 6195.84
Molecular Weight: 6200.04

**Ins-AC**(∇PheB1)

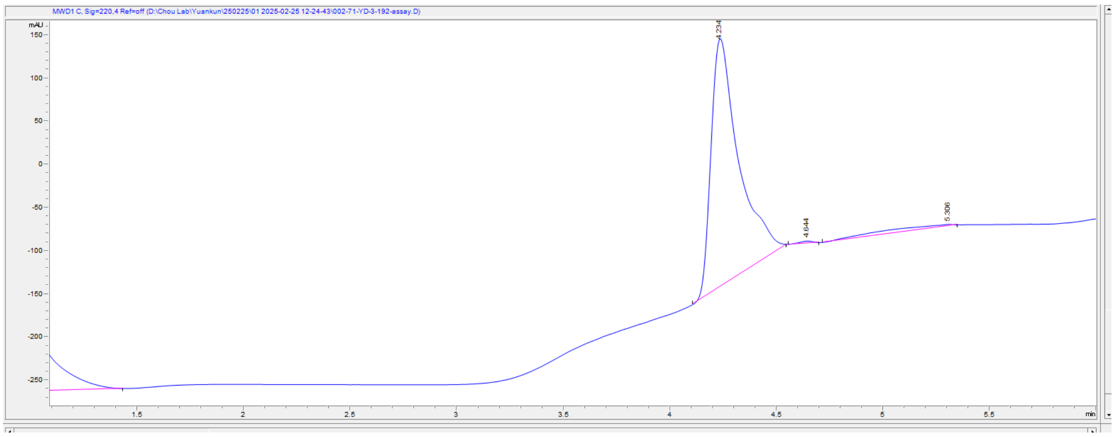

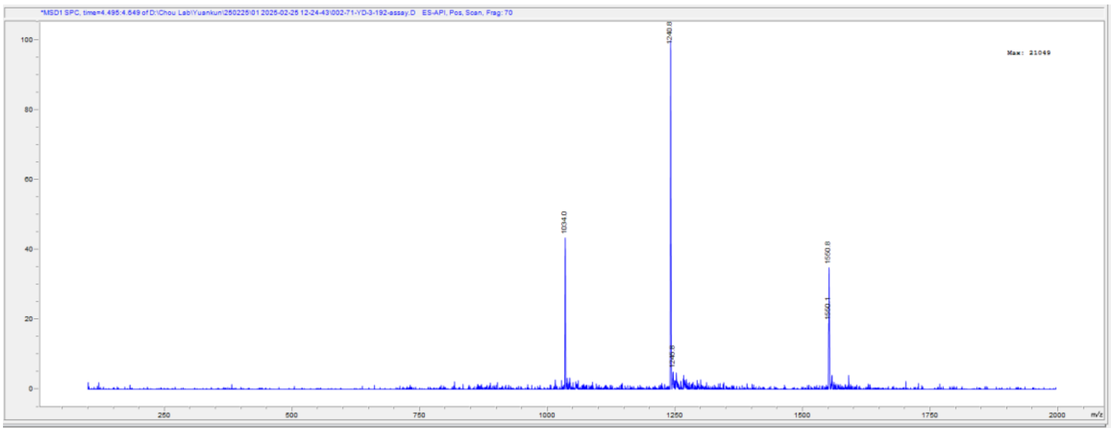


Top: UV traces of the purified **Ins-AC(▽PheB1)** at 220 nm using Phenomenex C18 column (Jupiter® 50 × 2 mm, 5 μm, 300 Å); Bottom: ESI-MS data of the purified **Ins-AC(▽PheB1)**. Calc. for $C_{271}H_{411}N_{71}O_{84}S_6$: 6200.04 Da (average isotopes), (m/z) $[M+4H]^{4+}$:1551.0, $[M+5H]^{5+}$:1241.0, $[M+6H]^{6+}$:1034.0; observed deconvoluted MS: 6199.25, $[M+4H]^{4+}$:1550.8, $[M+5H]^{5+}$:1240.8, $[M+6H]^{6+}$:1034.0.

***Ins-AC(ValB2Glu)***

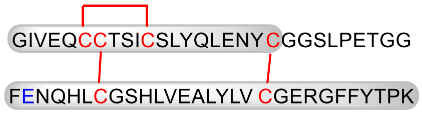


**Ins-AC**(ValB2Glu)

Chemical Formula: $C_{280}H_{417}N_{71}O_{88}S_6$
Exact Mass: 6373.87
Molecular Weight: 6378.19

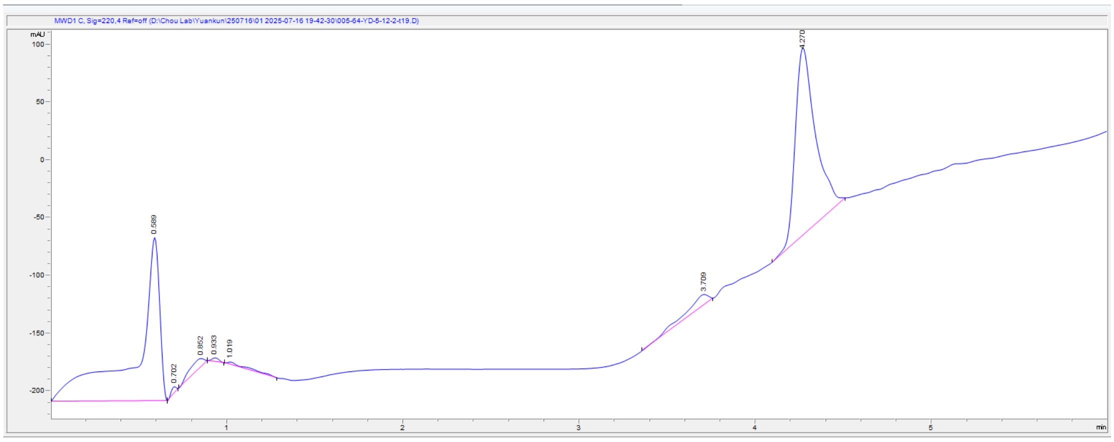


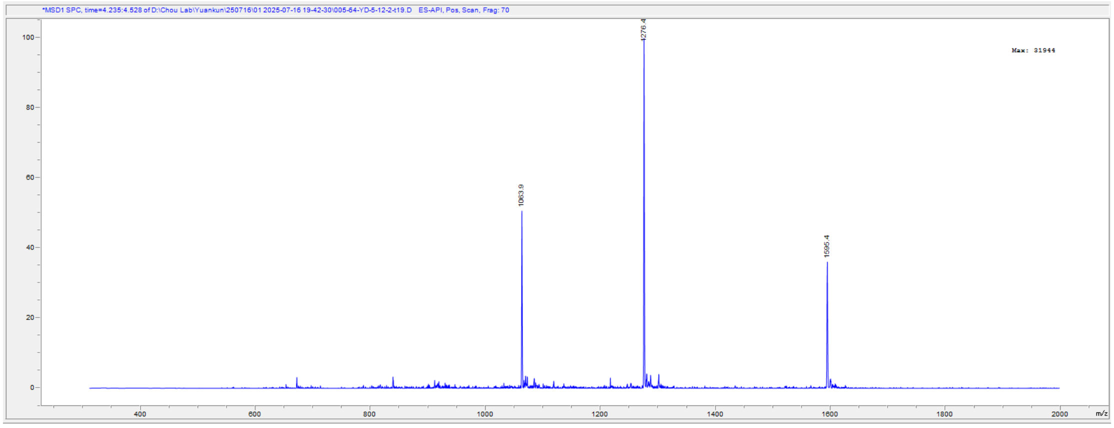


Top: UV traces of the purified **Ins-AC(ValB2Glu)** at 220 nm using Phenomenex C18

column (Jupiter® 50 × 2 mm, 5 µm, 300 Å); Bottom: ESI-MS data of the purified **Ins-AC(ValB2Glu)**. Calc. for $C_{280}H_{417}N_{71}O_{88}S_6$: 6378.19 Da (average isotopes), (m/z) $[M+4H]^{4+}$:1595.0, $[M+5H]^{5+}$:1276.2, $[M+6H]^{6+}$:1063.8; observed deconvoluted MS: 6377.35, $[M+4H]^{4+}$:1595.4, $[M+5H]^{5+}$:1276.4, $[M+6H]^{6+}$:1063.9.

***<u>Ins-AC(AsnB3Ala)</u>***

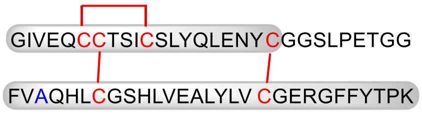


Chemical Formula: $C_{279}H_{418}N_{70}O_{85}S_6$
Exact Mass: 6300.89
Molecular Weight: 6305.18

**Ins-AC**(AsnB3Ala)**-AC**

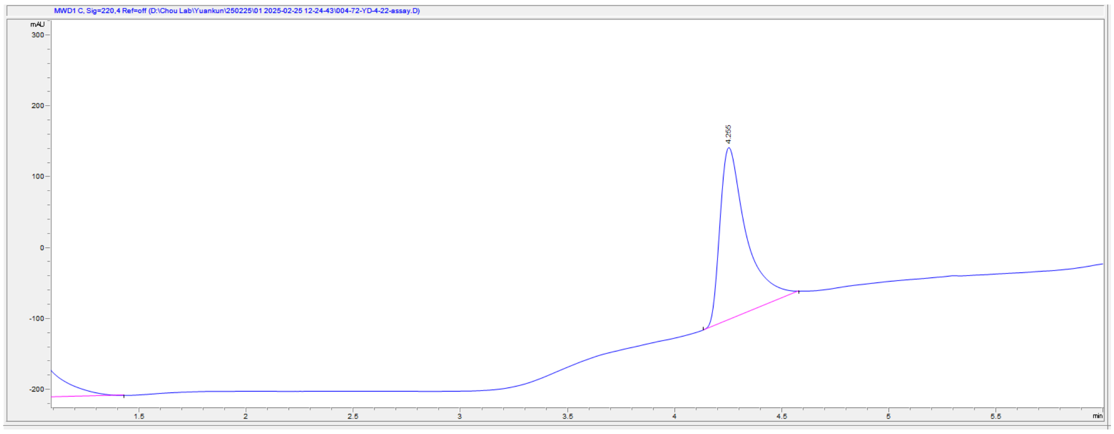


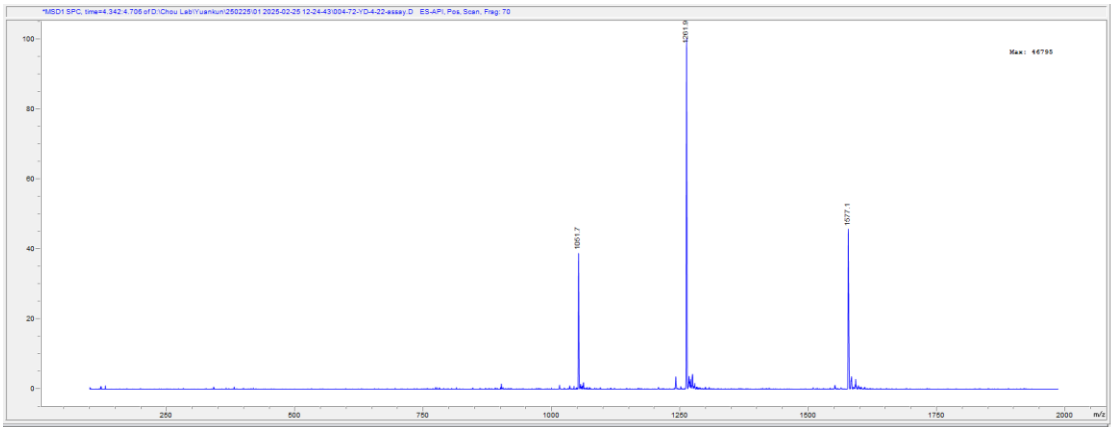


Top: UV traces of the purified **Ins-AC(AsnB3Ala)** at 220 nm using Phenomenex C18 column (Jupiter® 50 × 2 mm, 5 µm, 300 Å); Bottom: ESI-MS data of the purified **Ins-AC(AsnB3Ala)**. Calc. for $C_{279}H_{418}N_{70}O_{85}S_6$: 6305.18 Da (average isotopes), (m/z) $[M+4H]^{4+}$:1576.7, $[M+5H]^{5+}$:1261.6, $[M+6H]^{6+}$:1051.7; observed deconvoluted MS: 6304.34, $[M+4H]^{4+}$:1577.1, $[M+5H]^{5+}$:1261.9, $[M+6H]^{6+}$:1051.7.

***<u>Ins-AC(GlnB4Ala)</u>***

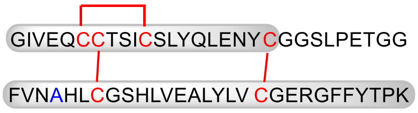


Chemical Formula: $C_{278}H_{416}N_{70}O_{85}S_6$
Exact Mass: 6286.87
Molecular Weight: 6291.15

**Ins-AC**(GlnB4Ala)

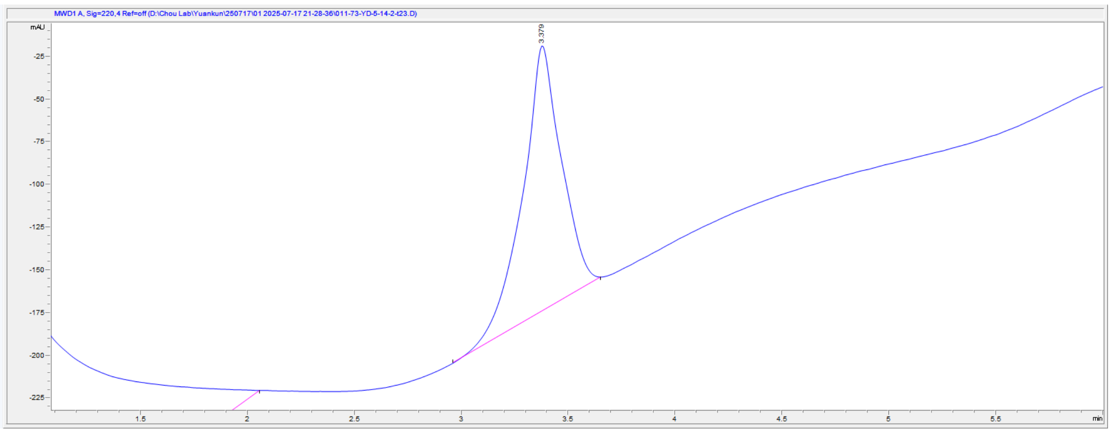


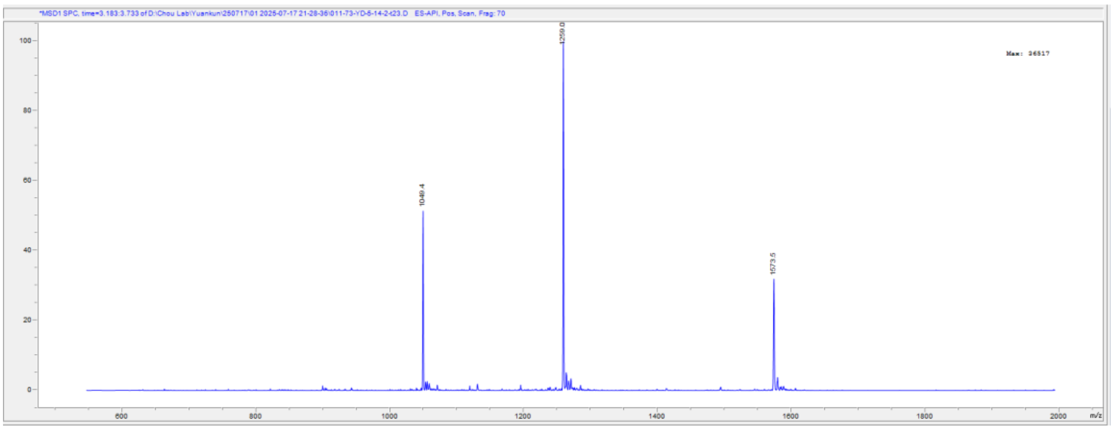


Top: UV traces of the purified **Ins-AC(GlnB4Ala)** at 220 nm using Phenomenex C18 column (Jupiter® 50 × 2 mm, 5 µm, 300 Å); Bottom: ESI-MS data of the purified **Ins-AC(GlnB4Ala)**. Calc. for $C_{278}H_{416}N_{70}O_{85}S_6$: 6291.15 Da (average isotopes), (m/z) $[M+4H]^{4+}$:1573.5, $[M+5H]^{5+}$:1259.2, $[M+6H]^{6+}$:1049.2; observed deconvoluted MS: 6290.36, $[M+4H]^{4+}$:1573.5, $[M+5H]^{5+}$:1259.0, $[M+6H]^{6+}$:1049.4.

***Ins-AC(HisB5Ala)***

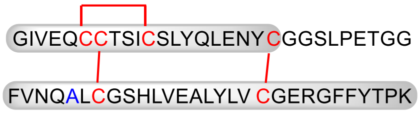


**Ins-AC**(HisB5Ala)

Chemical Formula: $C_{277}H_{417}N_{69}O_{86}S_6$
Exact Mass: 6277.87
Molecular Weight: 6282.14

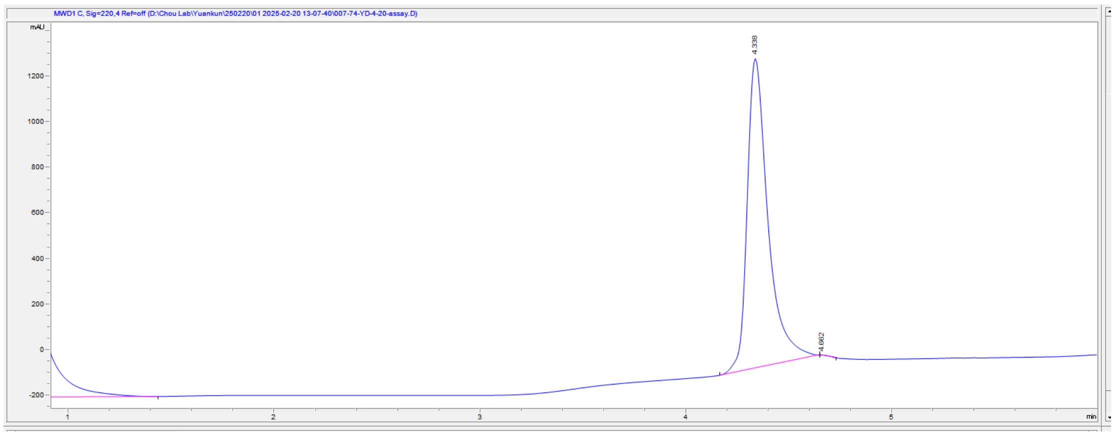

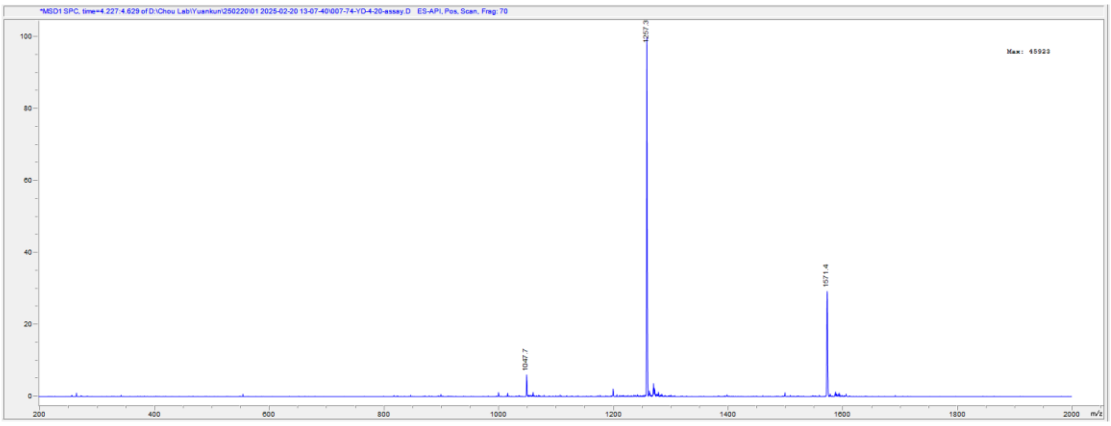


Top: UV traces of the purified **Ins-AC(HisB5Ala)** at 220 nm using Phenomenex C18 column (Jupiter® 50 × 2 mm, 5 μm, 300 Å); Bottom: ESI-MS data of the purified **Ins-AC(HisB5Ala)**. Calc. for $C_{277}H_{417}N_{69}O_{86}S_6$: 6282.14 Da (average isotopes), (m/z) $[M+4H]^{4+}$:1571.2, $[M+5H]^{5+}$:1257.4, $[M+6H]^{6+}$:1047.7; observed deconvoluted MS: 6281.27, $[M+4H]^{4+}$:1571.4, $[M+5H]^{5+}$:1257.3, $[M+6H]^{6+}$:1047.7.

***Ins-AC(SerA12Ala)***

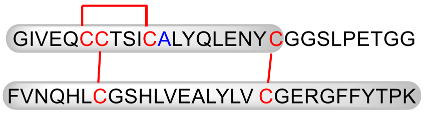


Chemical Formula: $C_{280}H_{419}N_{71}O_{85}S_6$
Exact Mass: 6327.90
Molecular Weight: 6332.20

**Ins-AC**(SerA12Ala)

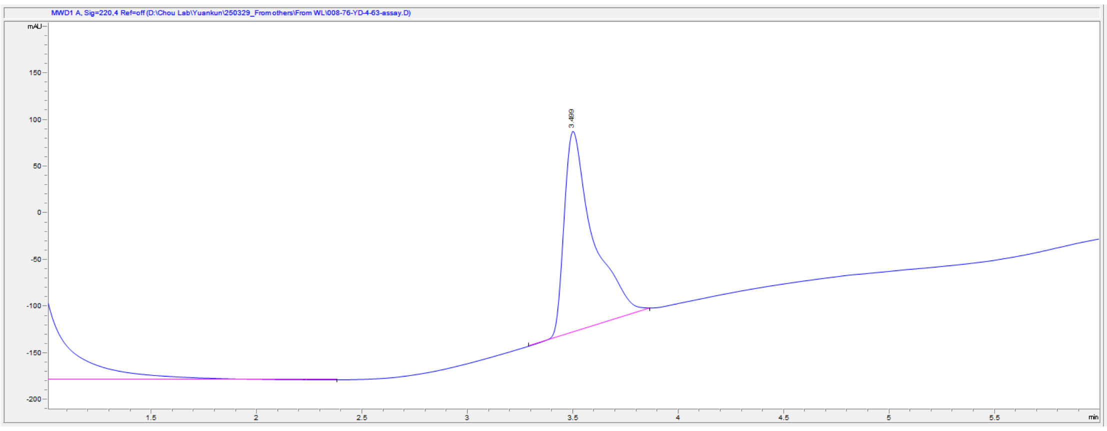


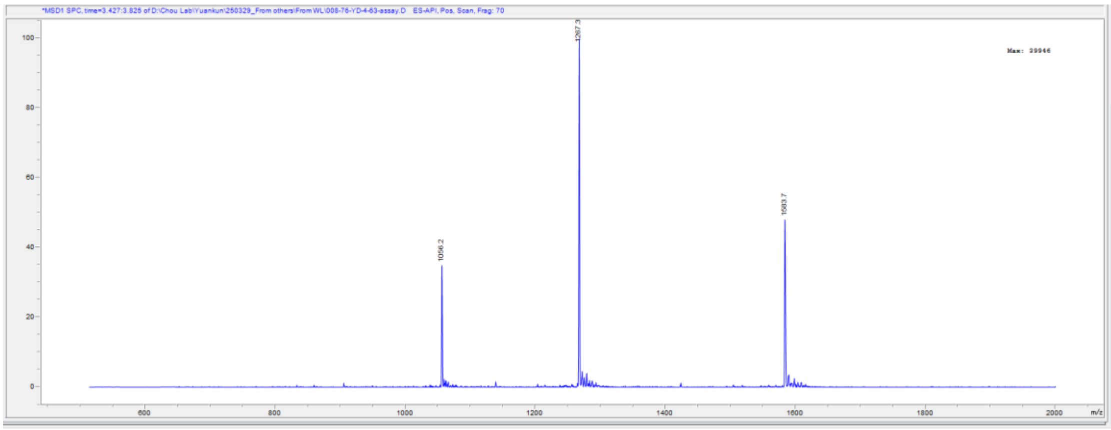


Top: UV traces of the purified **Ins-AC(SerA12Ala)** at 220 nm using Phenomenex C18

column (Jupiter® 50 × 2 mm, 5 μm, 300 Å); Bottom: ESI-MS data of the purified **Ins-AC(SerA12Ala)**. Calc. for $C_{280}H_{419}N_{71}O_{85}S_{6}$: 6332.20 Da (average isotopes), (m/z) $[M+4H]^{4+}$:1584.0, $[M+5H]^{5+}$:1267.0, $[M+6H]^{6+}$:1056.2; observed deconvoluted MS: 6331.32, $[M+4H]^{4+}$:1583.7, $[M+5H]^{5+}$:1267.3, $[M+6H]^{6+}$:1056.2.

#### ***Ins-AC-S2(▽PheB1)***

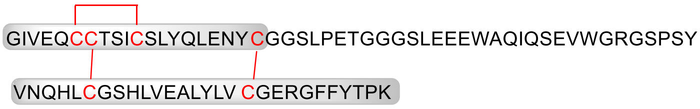


**Ins-AC-S2**(▽PheB1)

Chemical Formula: $C_{381}H_{567}N_{99}O_{122}S_{6}$
Exact Mass: 8672.95
Molecular Weight: 8678.66

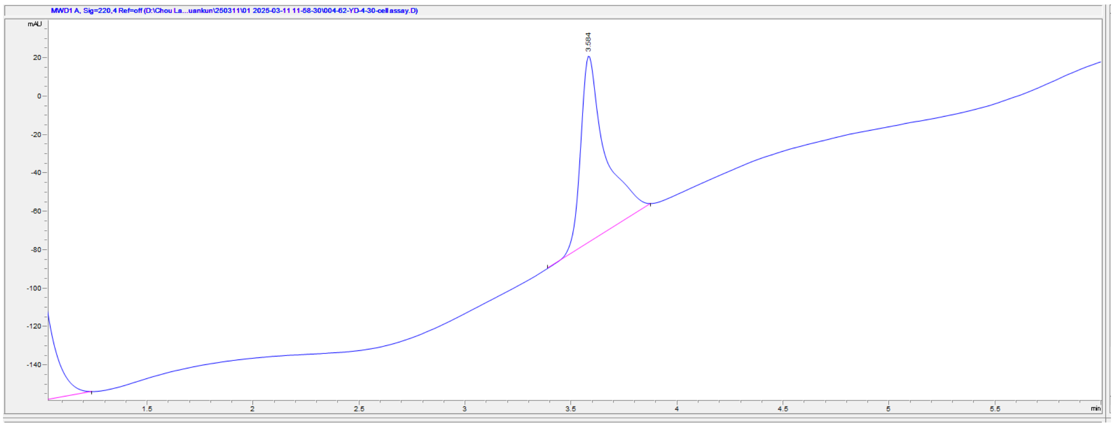


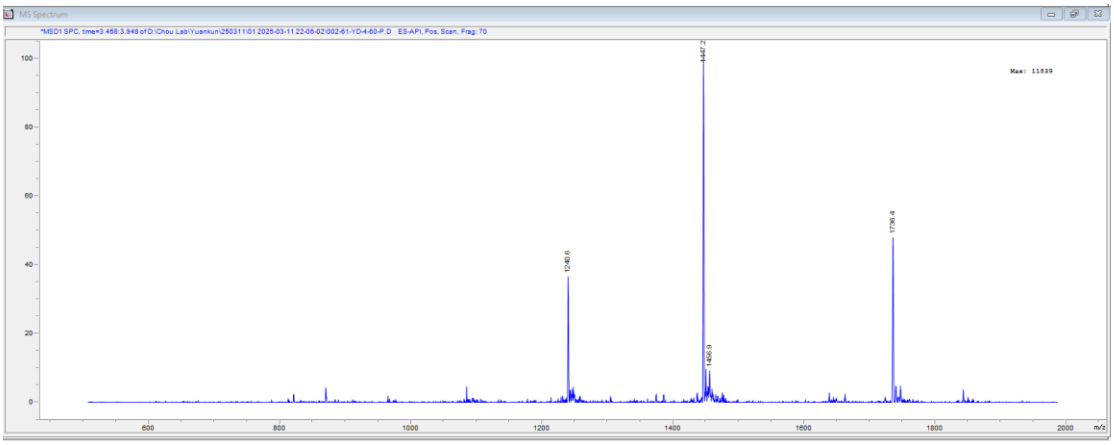


Top: UV traces of the purified **Ins-AC-S2(▽PheB1)** at 220 nm using Phenomenex C4 column (Jupiter® 50 × 2 mm, 5 μm, 300 Å); Bottom: ESI-MS data of the purified **Ins-AC-S2(▽PheB1)**. Calc. for $C_{381}H_{567}N_{99}O_{122}S_{6}$: 8678.66 Da (average isotopes), (m/z) $[M+5H]^{5+}$:1736.2, $[M+6H]^{6+}$:1447.0, $[M+7H]^{7+}$:1240.4; observed deconvoluted MS: 8677.99, $[M+5H]^{5+}$:1736.4, $[M+6H]^{6+}$:1447.2, $[M+7H]^{7+}$:1240.6.

#### ***Ins-AC-S2(ValB2Glu)***

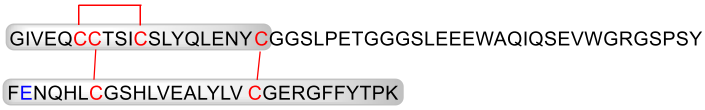


**Ins-AC-S2**(ValB2Glu)

Chemical Formula: $C_{390}H_{574}N_{100}O_{125}S_{6}$
Exact Mass: 8850.00
Molecular Weight: 8855.82

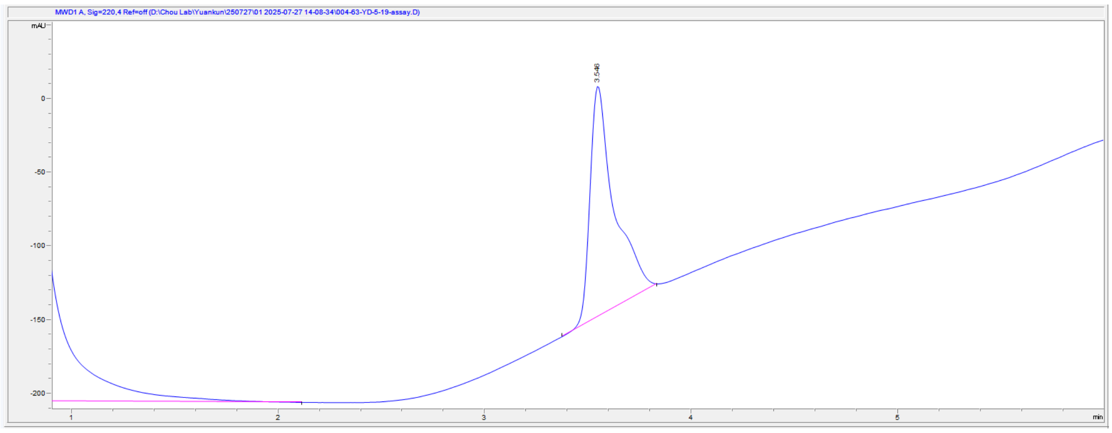

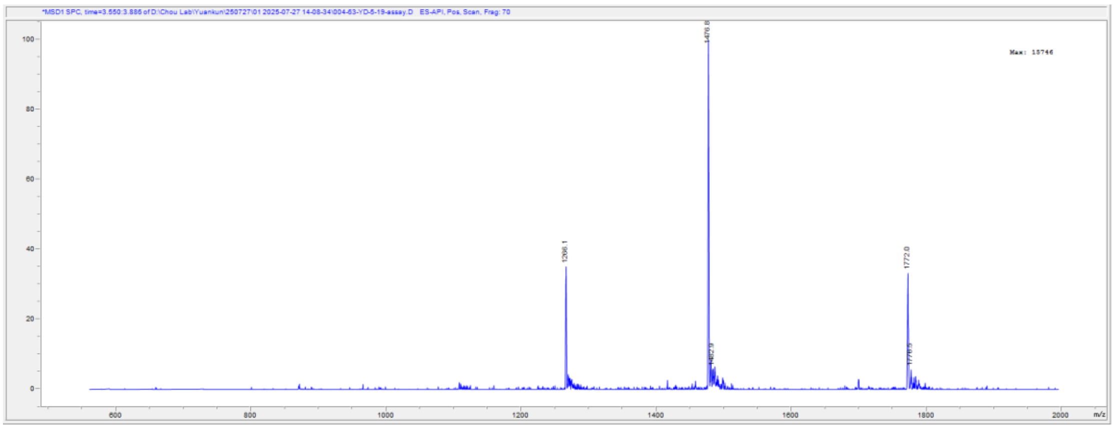

Top: UV traces of the purified **Ins-AC-S2(ValB2Glu)** at 220 nm using Phenomenex C4 column (Jupiter® 50 × 2 mm, 5 µm, 300 Å); Bottom: ESI-MS data of the purified **Ins-AC-S2(ValB2Glu)**. Calc. for $C_{390}H_{574}N_{100}O_{125}S_6$: 8855.82 Da (average isotopes), (m/z) $[M+5H]^{5+}$:1771.8, $[M+6H]^{6+}$:1476.7, $[M+7H]^{7+}$:1265.9; observed deconvoluted MS: 8854.84, $[M+5H]^{5+}$:1772.0, $[M+6H]^{6+}$:1476.8, $[M+7H]^{7+}$:1266.1.

#### ***Ins-AC-S2(AsnB3Ala)***

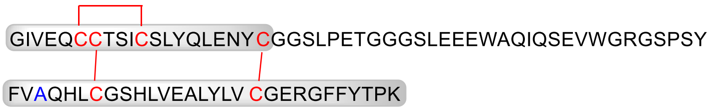


Chemical Formula: $C_{389}H_{575}N_{99}O_{122}S_6$
Exact Mass: 8777.02
Molecular Weight: 8782.81

**Ins-AC-S2**(AsnB3Ala)

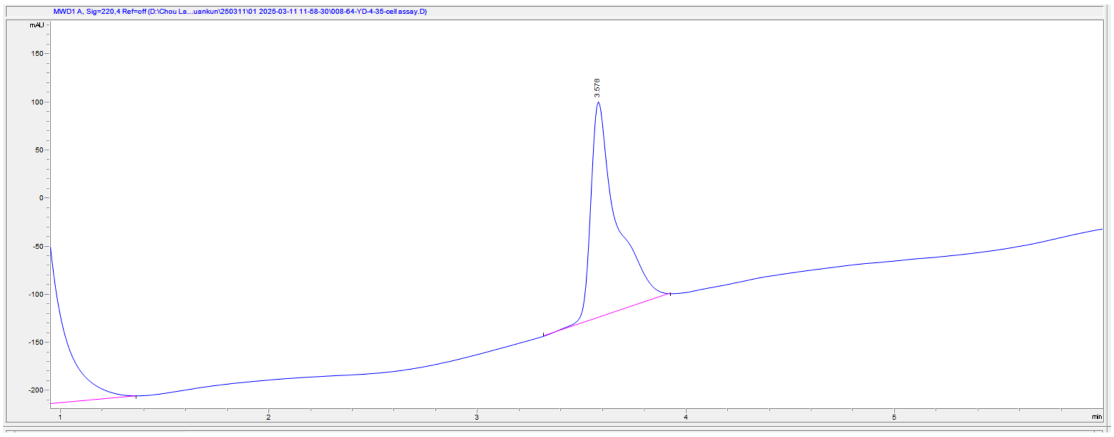

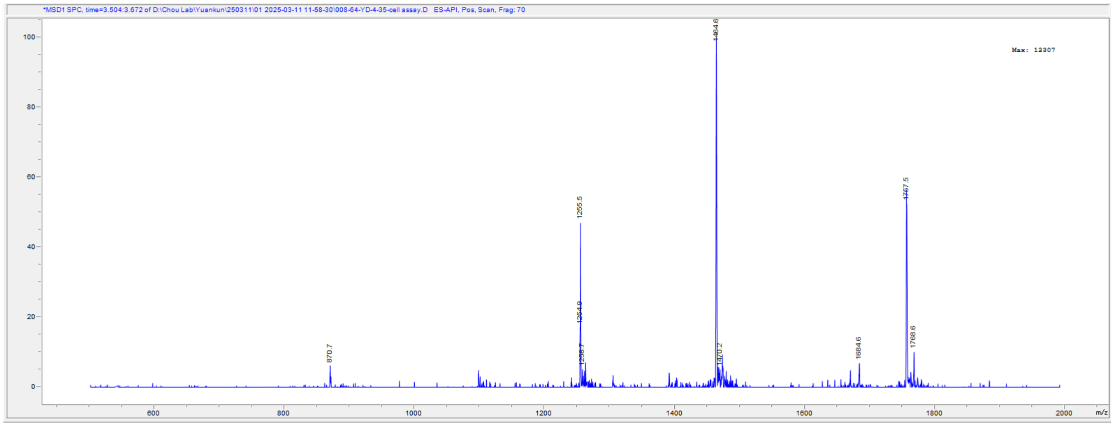


Top: UV traces of the purified **Ins-AC-S2(AsnB3Ala)** at 220 nm using Phenomenex C4 column (Jupiter® 50 × 2 mm, 5 μm, 300 Å); Bottom: ESI-MS data of the purified **Ins-AC-S2(AsnB3Ala)**. Calc. for $C_{389}H_{575}N_{99}O_{122}S_6$: 8782.81 Da (average isotopes), (m/z) $[M+5H]^{5+}$:1757.2, $[M+6H]^{6+}$:1464.5, $[M+7H]^{7+}$:1255.4; observed deconvoluted MS: 8782.26, $[M+5H]^{5+}$:1757.5, $[M+6H]^{6+}$:1464.6, $[M+7H]^{7+}$:1255.5.

#### ***Ins-AC-S2(GlnB4Ala)***

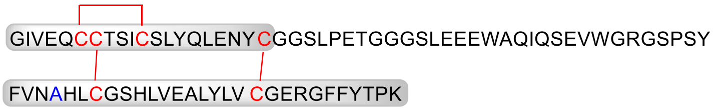


**Ins-AC-S2**(GlnB4Ala)

Chemical Formula: $C_{388}H_{573}N_{99}O_{122}S_6$
Exact Mass: 8763.00
Molecular Weight: 8768.78

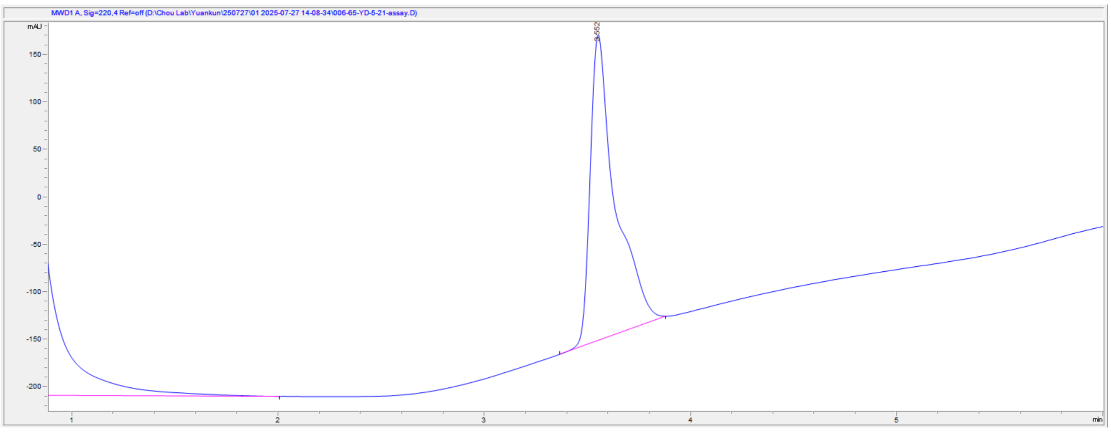


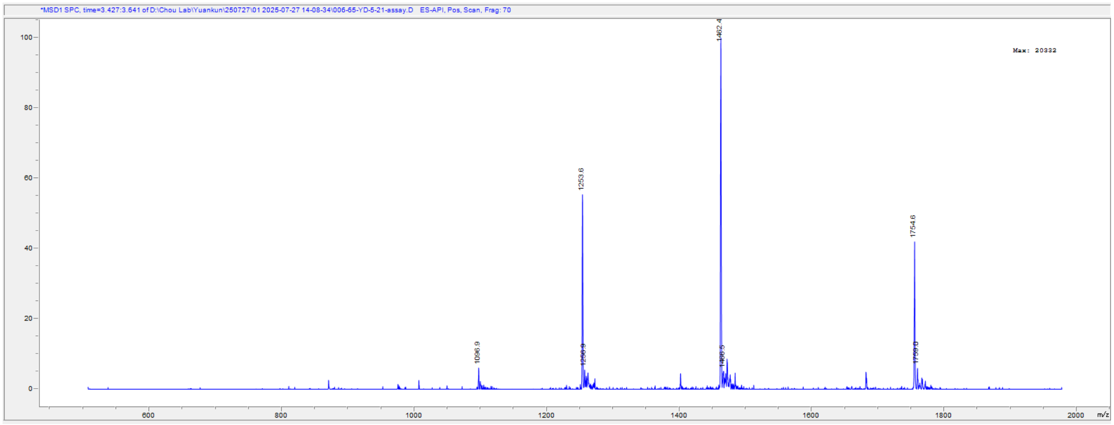


Top: UV traces of the purified **Ins-AC-S2(GlnB4Ala)** at 220 nm using Phenomenex

C4 column (Jupiter® 50 × 2 mm, 5 µm, 300 Å); Bottom: ESI-MS data of the purified **Ins-AC-S2(GlnB4Ala)**. Calc. for $C_{388}H_{573}N_{99}O_{122}S_6$: 8768.78 Da (average isotopes), (m/z) $[M+5H]^{5+}$:1754.4, $[M+6H]^{6+}$:1462.2, $[M+7H]^{7+}$:1253.4; observed deconvoluted MS: 8767.79, $[M+5H]^{5+}$:1754.6, $[M+6H]^{6+}$:1462.4, $[M+7H]^{7+}$:1253.6.

#### ***Ins-AC-S2(HisB5Ala)***

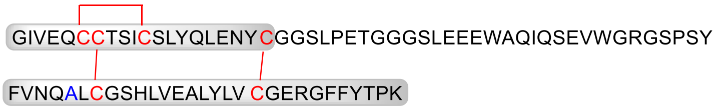


Chemical Formula: $C_{387}H_{574}N_{98}O_{123}S_6$
Exact Mass: 8754.00
Molecular Weight: 8759.77

**Ins-AC-S2**(HisB5Ala)

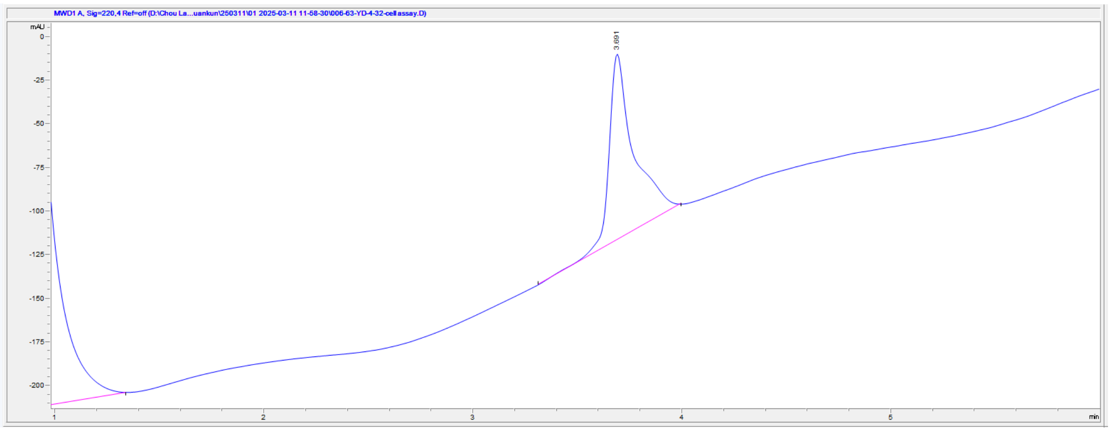


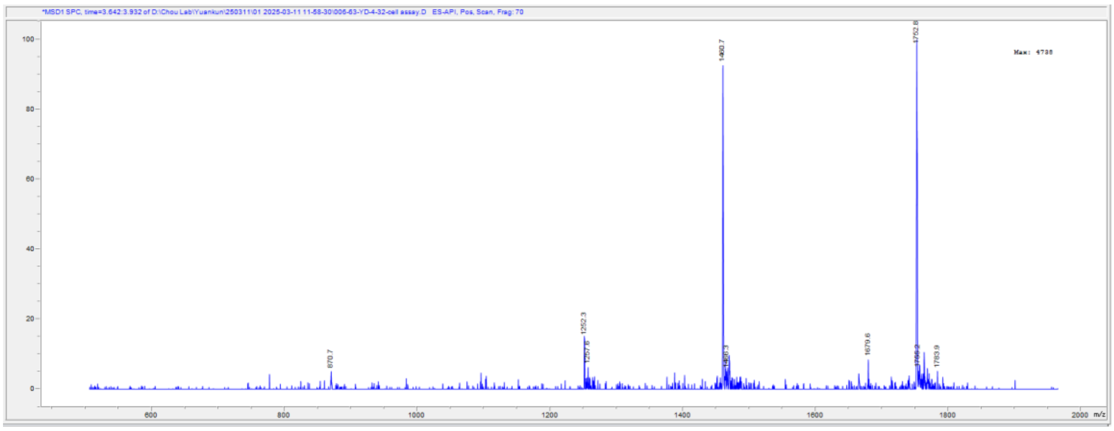


Top: UV traces of the purified **Ins-AC-S2(HisB5Ala)** at 220 nm using Phenomenex C4 column (Jupiter® 50 × 2 mm, 5 µm, 300 Å); Bottom: ESI-MS data of the purified **Ins-AC-S2(HisB5Ala)**. Calc. for $C_{387}H_{574}N_{98}O_{123}S_6$: 8759.77 Da (average isotopes), (m/z) $[M+5H]^{5+}$:1752.6, $[M+6H]^{6+}$:1460.7, $[M+7H]^{7+}$:1252.2; observed deconvoluted MS: 8759.07, $[M+5H]^{5+}$:1752.8, $[M+6H]^{6+}$:1460.7, $[M+7H]^{7+}$:1252.3.

#### ***Ins-AC-S2(SerA12Ala)***

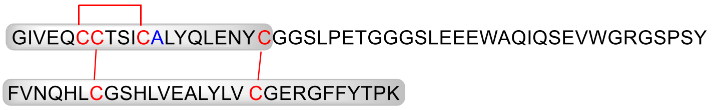


Chemical Formula: $C_{390}H_{576}N_{100}O_{122}S_6$
Exact Mass: 8804.03
Molecular Weight: 8809.84

**Ins-AC-S2**(SerA12Ala)

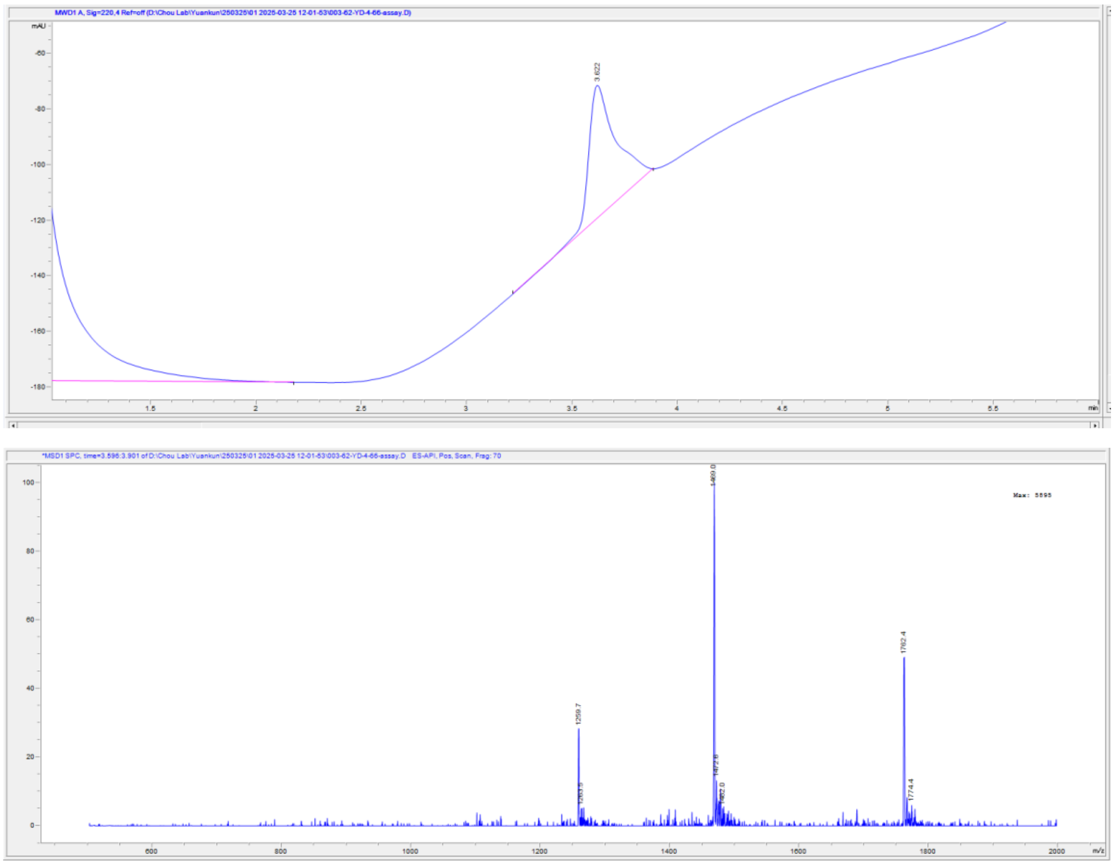

Top: UV traces of the purified **Ins-AC-S2(SerA12Ala)** at 220 nm using Phenomenex C4 column (Jupiter® 50 × 2 mm, 5 µm, 300 Å); Bottom: ESI-MS data of the purified **Ins-AC-S2(SerA12Ala)**. Calc. for $C_{390}H_{576}N_{100}O_{122}S_6$: 8809.84 Da (average isotopes), (m/z) $[M+5H]^{5+}$:1762.6, $[M+6H]^{6+}$:1469.0, $[M+7H]^{7+}$:1259.3; observed deconvoluted MS: 8808.94, $[M+5H]^{5+}$:1762.4, $[M+6H]^{6+}$:1469.0, $[M+7H]^{7+}$:1259.7.
